# Disentangled multi-subject and social behavioral representations through a constrained subspace variational autoencoder (CS-VAE)

**DOI:** 10.1101/2022.09.01.506091

**Authors:** Daiyao Yi, Simon Musall, Anne Churchland, Nancy Padilla-Coreano, Shreya Saxena

**Author notes:** {, }.

## Abstract

Effectively modeling and quantifying behavior is essential for our understanding of the brain. Modeling behavior in naturalistic settings in social and multi-subject tasks remains a significant challenge. Modeling the behavior of different subjects performing the same task requires partitioning the behavioral data into features that are common across subjects, and others that are distinct to each subject. Modeling social interactions between multiple individuals in a freely-moving setting requires disentangling effects due to the individual as compared to social investigations. To achieve flexible disentanglement of behavior into interpretable latent variables with individual and across-subject or social components, we build on a semi-supervised approach to partition the behavioral subspace, and propose a novel regularization based on the Cauchy-Schwarz divergence to the model. Our model, known as the constrained subspace variational autoencoder (CS-VAE), successfully models distinct features of the behavioral videos across subjects, as well as continuously varying differences in social behavior. Our approach vastly facilitates the analysis of the resulting latent variables in downstream tasks such as uncovering disentangled behavioral motifs, the efficient decoding of a novel subject’s behavior, and provides an understanding of how similarly different animals perform innate behaviors.

## 1 Introduction

Effective study of the relationship between neural signals and ensuing behavior relies on our ability to measure and adequately quantify behavior. Historically, behavior has been quantified by a very small number of markers as the subject performs the task, for example, force sensors on levers. However, advancement in hardware and storage capabilities, as well as computational methods applied to video data, has allowed us to increase the quality and capability of behavioral recordings to videos of the entire subject that can be processed and analyzed quickly. It is now widely recognized that understanding the relationship between complex neural activity and high-dimensional behavior is a major step in understanding the brain that has been undervalued in the past [1, 2]. However, the analysis of high-dimensional behavioral video data across subjects is still a nascent field, due to the lack of adequate tools to efficiently disentangle behavioral features related to different subjects. Moreover, as recording modalities become light-weight and portable, neural and behavioral recordings can be performed in more naturalistic settings, which are difficult for behavioral analysis tools to disentangle due to changing scenes.

Although pose estimation tools that track various body parts in a behavioral video are very popular, they fail to capture smaller movements and rely on the labeler to judge which parts of the scene are important to track [3, 4, 5, 6, 7]. Unsupervised techniques have gained traction to circumvent these problems. These include directly applying dimensionality reduction methods such as Principal Component Analysis (PCA) and Variational Autoencoders (VAEs) to video data [2, 8, 9]. However, understanding or segmentation of the latent variables is difficult for any downstream tasks such as motif generation. To combine the best of both worlds, semi-supervised VAEs have been used for the joint estimation of tracked body parts and unsupervised latents that can effectively describe the entire image [2]. These have not been applied to across-subject data, with the exception of [10], where the authors directly use a frame of each subject’s video as a context frame to define individual differences; however, this method only works with a *discrete* set of *labeled* sessions or subjects. These methods fail when applied without labeled subject data, or more importantly, when analyzing freely-behaving social behavior, due to continuously shifting image distributions that confound the latent space.

With increasing capabilities to effectively record more naturalistic data in neuroscience, there is a growing demand for behavioral analysis methods that are tailored to these settings. In this work, we model a continuously varying distribution of images, such as in freely moving and multi-subject behavior, by using a novel loss term called the Cauchy-Schwarz Divergence (CSD) [11, 12]. By applying the CSD loss term, a subset of the latents can be automatically projected on a pre-defined and flexible distribution, thus leading to an unbiased approach towards latent separation. Here, the CSD is an effective variational regularizer that separates the latents corresponding to images with different appearances, thus successfully capturing ‘background’ information of an individual. This background information can be the difference in lighting during the experiment, the difference in appearance across mice in a multi-subject dataset, or the presence of another subject in the same field of view as in a social interaction dataset.

To further demonstrate the utility of our approach, we show that we can recover behavioral motifs from the resulting latents in a seamless manner. We recover (a) the same motifs across different animals performing the same task, and (b) motifs pertaining to social interactions in a freely moving task with two animals. Furthermore, we show the neural decoding of multiple animals in a unified model, with benefits towards the efficient decoding of the behavior of a novel subject. Finally, we compare the commonalities in neural activity across different trials in the same subject to those across subjects for different types of behavior motifs, e.g. task-related and spontaneous.

### Related Works

Pose estimation tools such as DeepLabCut (DLC) and LEAP have been broadly applied to neuroscience experiments to track the body parts of animals performing different tasks, including in the social setting [3, 4, 5, 6, 7]. These are typically supervised techniques that require extensive manual labeling. Although these methods can be sample-efficient due to the use of transfer learning methods, they still depend inherently on the quality of the manual labels, which can differ across labelers. Moreover, these methods may be missing key information in these behavioral videos that are not captured by tracking the body parts, for example, movements of the face, the whiskers, and smaller muscles that comprise a subject’s movements.

Emerging unsupervised methods have demonstrated significant potential in directly modeling behavioral videos. A pioneer in this endeavor was MoSeq, a behavioral video analysis tool that encodes high dimensional behavior by directly applying PCA to the data [13, 9]. Behavenet is similar to MoSeq, but uses autoencoders to more effectively reduce the dimensionality of the representation [8]. However, the corresponding latent variables in these models are typically not interpretable. To add interpretability, the Partitioned Subspace VAE (PS-VAE) [2] formulates a semi-supervised approach that uses the labels generated using pose estimation methods such as DLC in order to partition the latent representation into both supervised and unsupervised subspaces. The ‘supervised’ latent subspace captures the parts that are labeled by pose estimation software, while the ‘unsupervised’ latent subspace encodes the parts of the image that have not been accounted for by the supervised space. While PS-VAE is very effective for a single subject, it does not address latent disentaglement in the ‘unsupervised’ latent space, and is not able to model multi-subject or social behavioral data.

Modeling multiple sessions has recently been examined in two approaches: MSPS-VAE and DBE [2, 10]. Both of these are confined to modeling head-fixed animals with a pre-specified number of sessions or subjects. In MSPS-VAE, an extension to PS-VAE, a latent subspace is introduced in the model that encodes the static differences across sessions. In DBE, a context frame from each session or subject is used as a static input to generate the behavioral embeddings. Two notable requirements of applying both these methods is the presence of a discrete number of labeled sessions or subjects in the dataset. Therefore, these are not well suited for naturalistic settings where the session / subject identity might not be known a priori, or the scene might be continuously varying, for example, in the case of subjects roaming in an open-field.

## 2 Results

### 2.1 CS-VAE Model Structure

Although existing pose estimation methods are capable enough to capture the body position of the animals in both open and contained space, tracking specific actions such as shaking and wriggling still remains a problem. However, a purely unsupervised or semi-supervised model such as a VAE or PS-VAE lacks the ability to extract meaningful and interoperable behaviors from multi-subject or social behavioral videos. One possible solution is to add another set of latent which could capture the variance across individuals and during social interactions. Instead of constraining the data points from different sessions or subjects to distinct parts of the subspace as in [2, 10], we directly constrain the latent subspace to a flexible prior distribution using a Cauchy-Schwarz regularizer as detailed in the Methods section. Ideally, this constrained subspace (CS) captures the difference between different animals in the case of a multi-subject task and the social interactions in a freely-behaving setting, while the supervised and unsupervised latents are free to capture the variables corresponding to the individual. The model structure described above is shown in Fig. 1. After the input frames go through a series of convolutional layers, the resulting latent splits into three sets. The first set contains the supervised latents, which encodes the specific body position as tracked by supervised tracking methods such as DLC. The unsupervised latents capture the rest of the individual’s behavior that are not captured by supervised latents. The CS latents capture the continuous difference across frames. The prior distribution can be changed to fit different experimental settings (and can be modeled as a discretized state space if so desired, making it close to the MSPS-VAE discussed in the Introduction).

**Figure 1:**
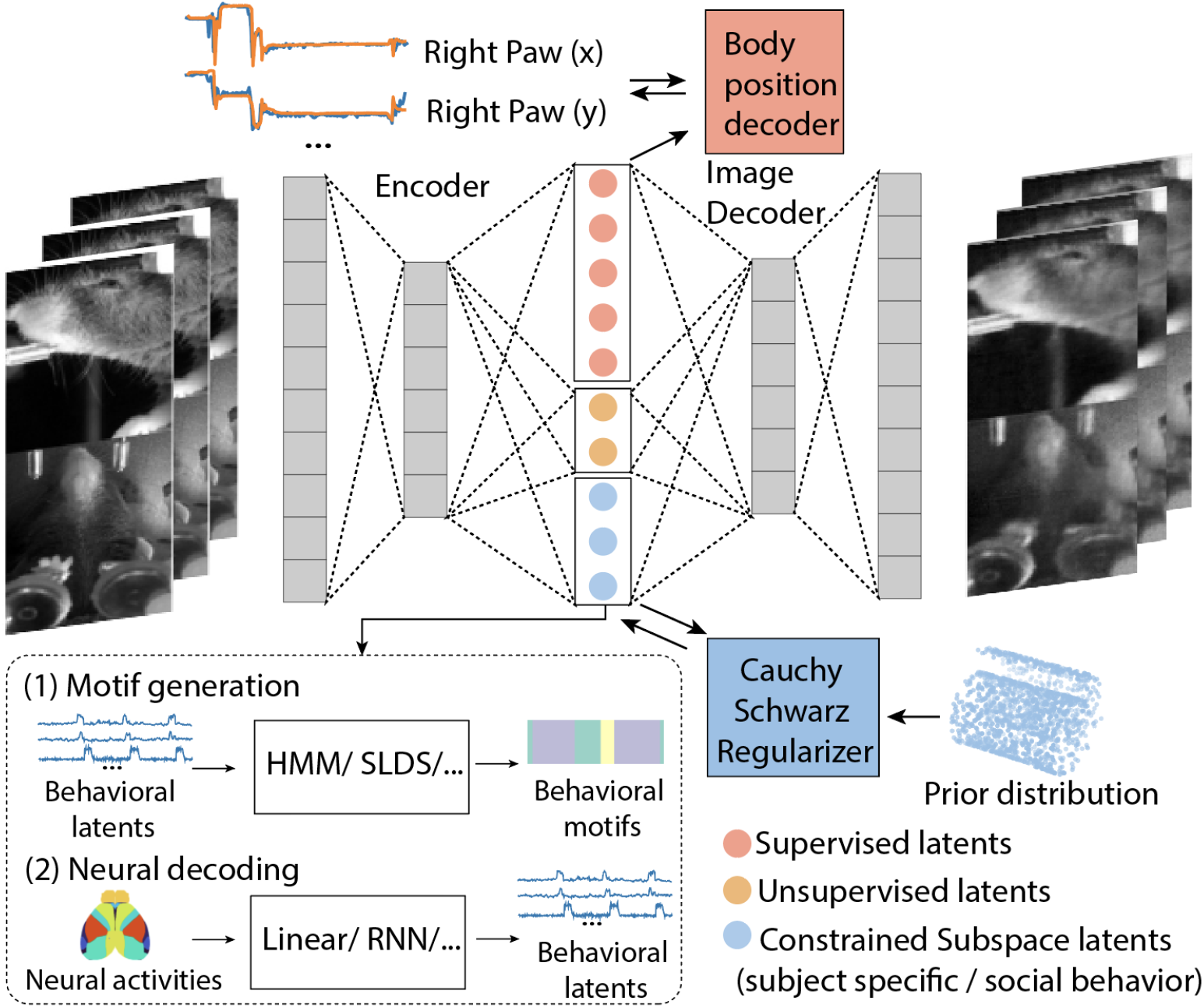
Overview of the Constrained Subspace Variational Autoencoder (CS-VAE). The latent space is divided in three parts: (1) the supervised latents decode the labeled body positions, (2) the unsupervised latents model the individual’s behavior that is not explained by the supervised latents, and (3) the constrained subspace latents model the continuously varying features of the image, e.g., relating to multi-subject or social behavior. After training the network, the generated latents can be applied to several downstream tasks. Here we show two example tasks: (1) Motif generation: we apply state space models such as hidden Markov models (HMM) and switched linear dynamical systems (SLDS), with the behavioral latent variables as the observations; (2) Neural decoding: with neural recordings such as widefield calcium imaging, corresponding behaviors can be efficiently predicted for novel subjects.

### 2.2 Modeling Smooth Variations in a Simulated Dataset

We performed a simulation study on the behavioral videos of one of the mice in the ‘Multi-Subject Behavior’ dataset detailed in Appendix .1. We applied a continuously varying contrast ratio throughout the trials (Fig. 2A) to model smoothly varying lighting differences across the dataset. We then randomly shuffled all the trials and trained a CS-VAE model with a swiss roll as a prior distribution. Here, the *R*^2^ for the supervised labels was 0.881 ± 0.05 (Fig. 2C), and the mean squared error (MSE) for reconstructing the entire frame was 0.0067 ± 0.0003, showing that both the images and the labels were fit well.

**Figure 2:**
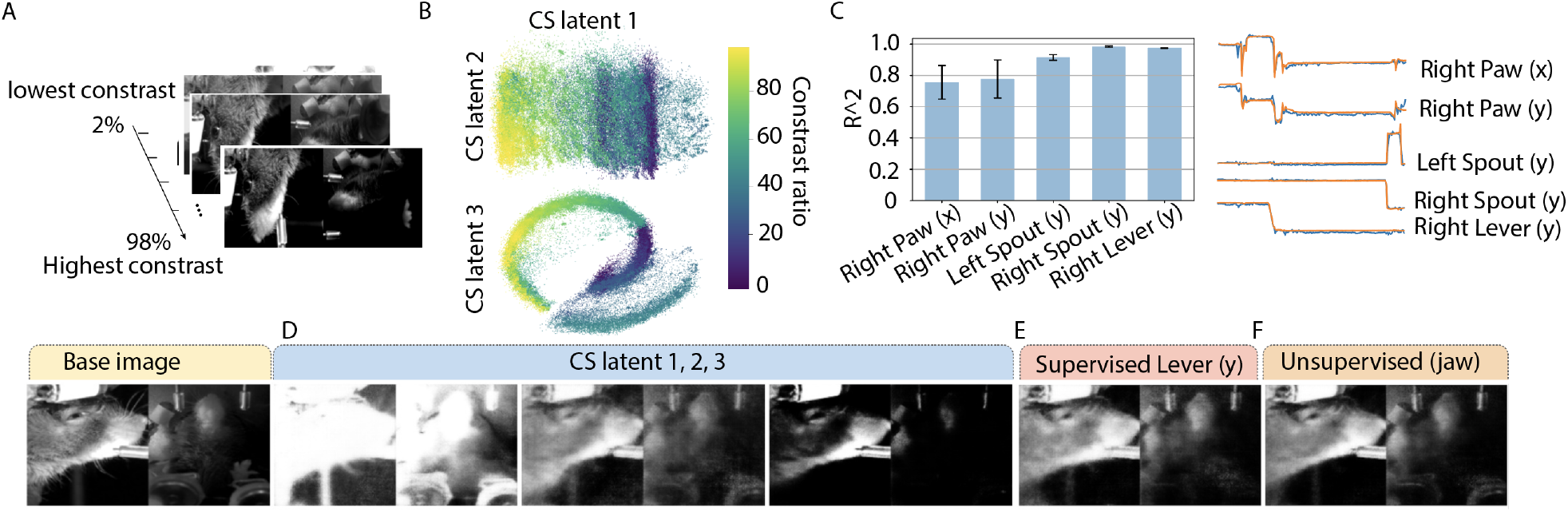
(A) Simulated dataset: behavioral videos from one mouse with artificially simulated differences in contrast. (B) Distribution occupied by the 3 CS latents.The constrained latents are distributed according to the pre-defined prior: a Swiss roll distribution. Different contrast ratios separate well in space. (C) Left: *R*^2^ values for label reconstruction; Right: visualization of label reconstruction for an example trial. Latent traversals for (D) CS latents, each of which captures lower, medium, and higher contrast rate. (E) An example supervised latent captures lever movement, and (F) an example unsupervised latent which captures jaw movement.

We show the CS latents recovered by the model in Fig. 2B, which follow the contrast ration distribution. We also show latent traversals in Fig. 2D-F, which demonstrate that the CS latent successfully captured the contrast changes in the frames (Fig. 2D), the supervised latent successfully captured the corresponding labeled body part (Fig. 2E), and the unsupervised latent captured parts of the individual’s body movement with a strong emphasis on the jaw (Fig. 2F). Thus, we show that smoothly varying changes in the videos are well captured by our model.

### 2.3 Modeling Multi-Subject Behavior

In a multi-subject behavioral task, we would like to disentangle the commonalities in behavior from the differences across subjects. Here, we test the CS-VAE on an experimental dataset with four different mice performing a two-alternative forced choice task (2AFC): head-fixed mice performed a self-initiated visual discrimination task, while the behavior was recorded from two different views (face and body). The behavioral video includes the head-fixed mice as well as experimental equipment such as the levers and the spouts. We labeled the right paw, the spouts, and the levers using DLC [3]. Neural activity in the form of widefield calcium imaging across the entire mouse dorsal cortex was simultaneously recorded with the behavior. The recording and preprocessing details are in [14, 15], and the preprocessing steps for the neural data are detailed in [15].

#### Reconstruction Accuracy

The CS-VAE model results in a mean label reconstruction accuracy *R*^2^ = 0.926 ± 0.02 (Fig. 3B,C), with the MSE for frame reconstruction as 0.00232 ± 7.7 · 10^*−*5^ (Fig. 3A). This was comparable to the results obtained using a PS-VAE model (*R*^2^ = 0.99 ± 0.004, MSE = 0.13 ± 4.5 · 10^*−*7^).

**Figure 3:**
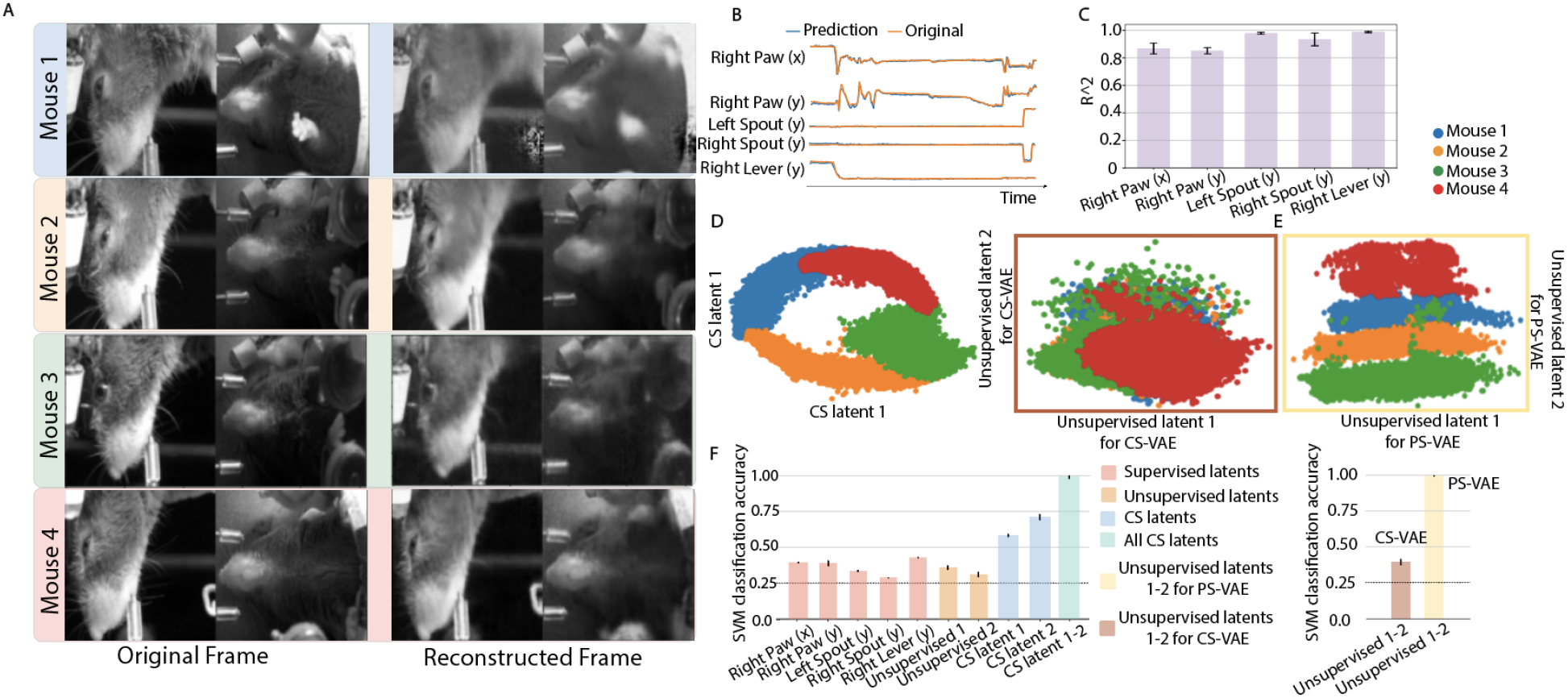
Modeling the behavior of four different mice. A. Image reconstruction result for an example frame from each mouse. B. Label reconstruction result for an example trial. C. *R*^2^ value for label reconstruction for all mice. D. (Left) CS latent and (Right) unsupervised latent distributions for all mice generated using our CS-VAE model. On the left, we see that the CS latent distribution follows the pre-defined prior distribution and is well separated; on the right, we see that the unsupervised latent distribution is well overlapped across mice. E. Unsupervised latent distribution for all mice generated using the comparison PS-VAE model, where the latents from different mice are separate from each other. F. SVM classification accuracy for classifying different mice using the CS-VAE and PS-VAE latents. The unsupervised latents generated by the CS-VAE has low classification accuracy, indicating across-subject representations, and the CS latents have a classification accuracy close to one, indicating good separation.

#### Disentangled Latent Space Representation

We show latent traversals for each mouse in Fig. 4, with the base image chosen separately for each mouse (videos in Supplementary Material 3). We see that, even for different mice, the supervised latent can successfully capture the corresponding labeled body part (Fig. 4A). The example unsupervised latent is shown to capture parts of the jaw of each mouse (Fig. 4B), and is well-localized, comparable with the example supervised latent. The CS latent dimension encodes many different parts of the image, and has a large effect on the appearance of the mouse, effectively changing the appearance from one mouse to another, signifying that it is useful in the case of modeling mouse-specific differences (Fig. 4C). We demonstrate the abilities of the CS latent in capturing the appearance of the mouse by directly changing the CS latent from one part of subspace to another (Figure 4D). The changes in appearance along with the invariance in actions shows the intraoperability between mice by only changing the CS latents in this model (Fig. 4D).

**Figure 4:**
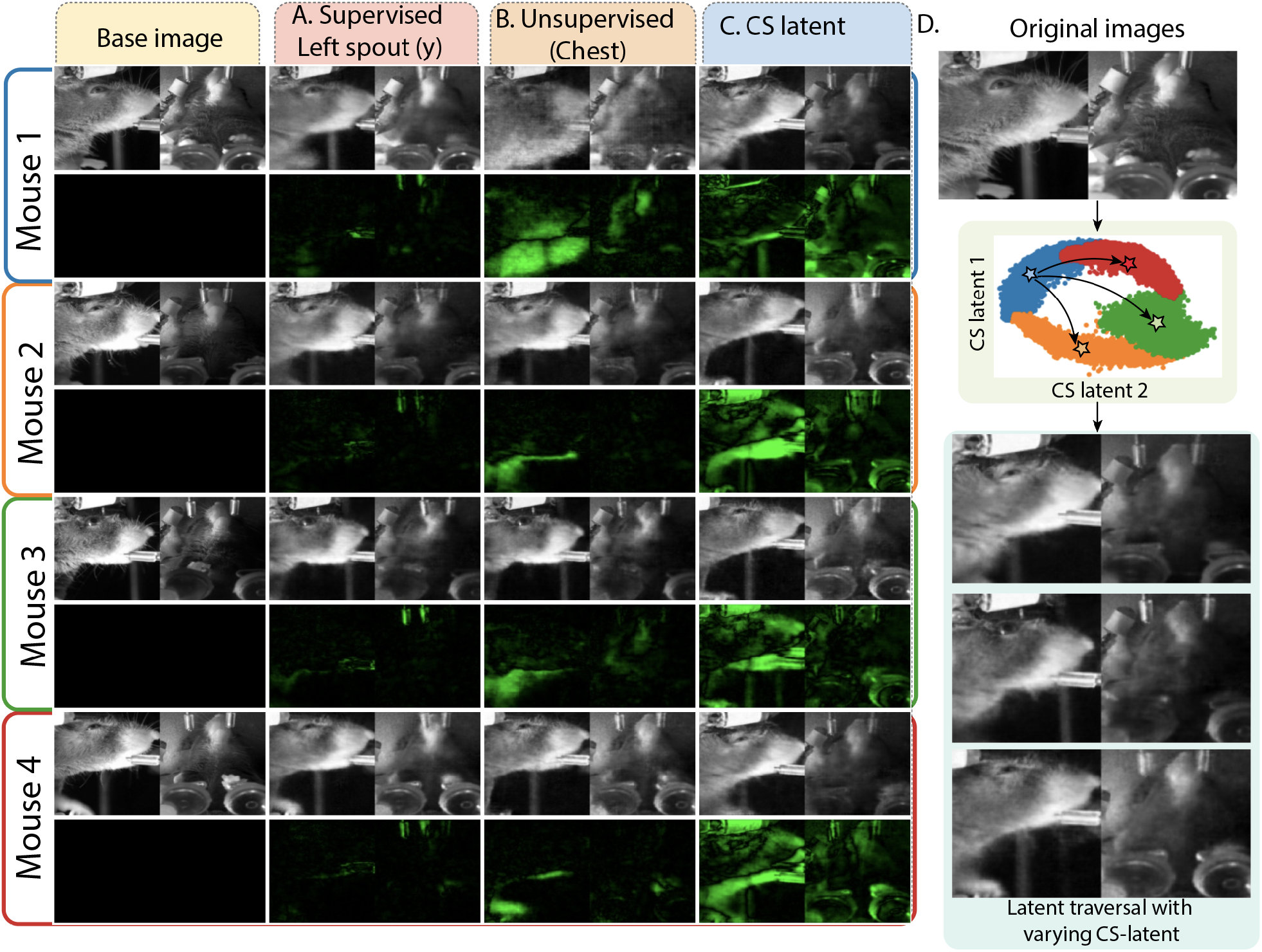
Latent traversals for behavioral modeling of four different mice for A. an example supervised latent that captures the left spout across all the subjects, B. an example unsupervised latent that captures the chest of the mice, and C. an example CS latent that successfully captures the mouse appearance. D. Changing the value of the CS latent in an example frame leads to a change in subject, while keeping the same action as in the example frame.

Ideally, we would like to uncover common across-subject variables using the supervised and unsupervised latents subspaces, and have the individual differences across subjects be encoded in the CS latents. Thus, we expect the unsupervised latents to not be able to classify the individual well. In fact, Fig. 3D,F show that the unsupervised latents overlap well across the four mice and perform close to chance level (0.25) in a subject-classification task using SVM (details in Appendix **??**). This signifies that unsupervised latents occupy the same values across all four mice and thus effectively capture across-subject behavior. In fact, we tested our latent space by choosing the same base image across the four mice, and found that the supervised and unsupervised latents from different mice can be used interchangeably to change the actions in the videos, also showing interoperability between different mice in these latent subspaces (Appendix .9).

This is in stark contrast to the CS latents, which are well separated across mice and are able to be classified well (Fig. 3D,F); thus, they effectively encode for individual differences across subjects. Note that our method did not *a prior* know the identity of the subjects, and thus this shows that the CS latents achieve separation in an unsupervised manner. We also note that the CS latents are distributed in the shape of the chosen prior distribution (a circle). The separation in the unsupervised latent space obtained by the baseline PS-VAE shown in Fig. 3E and the latents’ ability to classify different subjects (Fig. 3F) further validates the utility of CS-VAE.

Lastly, we trained the model while using prior distributions of different types, to understand the effect on the separability of the resulting latents. The separability was comparable across a number of different prior distributions, such as a swiss roll and a plane, signifying that the exact type of prior distribution does not play a large role.

#### Across-Subject Motif Generation

To further show that the supervised and unsupervised latents produced by CS-VAE are interoperable between the different mice, we apply a standard SLDS model (Appendix .6) to uncover the motifs using this across-subject subspace. As seen in the ethograms (left) and the histograms (right) in Fig. 5, the SLDS using the CS-VAE latents captures common states across different subjects, indicating that the latents are well overlapped across mice. The supervised latents related to equipment in the experiment, here the spout and lever, split the videos into four states (different colors in the ethograms in Fig. 5A), that we could independently match with ground truth obtained from sensors in these equipment. The histograms show that, as expected, these states occur with a very similar frequency across mice. We also explored the behavioral states related to the right paw. The resulting three states captured the idle vs. slightly moving vs. dramatically moving paw (Fig. 5B). The histograms show that these states also occur with a very similar frequency across mice. Videos for all these states are available in Supplementary Material 2. The inference drawn from supervised latents is directly proportional to the DLC labels. Hence, a similar conclusion can be arrived at by utilizing the DLC pose estimations. Nonetheless, the subsequent outcomes cannot be attained solely based on the poses. We extracted the behavioral states related to the unsupervised latents, which yielded 3 states related to raising of the paws (including grooming) and jaw movements (including licking) that are present in all four mice, as shown in Fig. 5C. We see that different mice have different tendencies to lick and groom, e.g., mouse 1 and 4 seem to groom more often.

**Figure 5:**
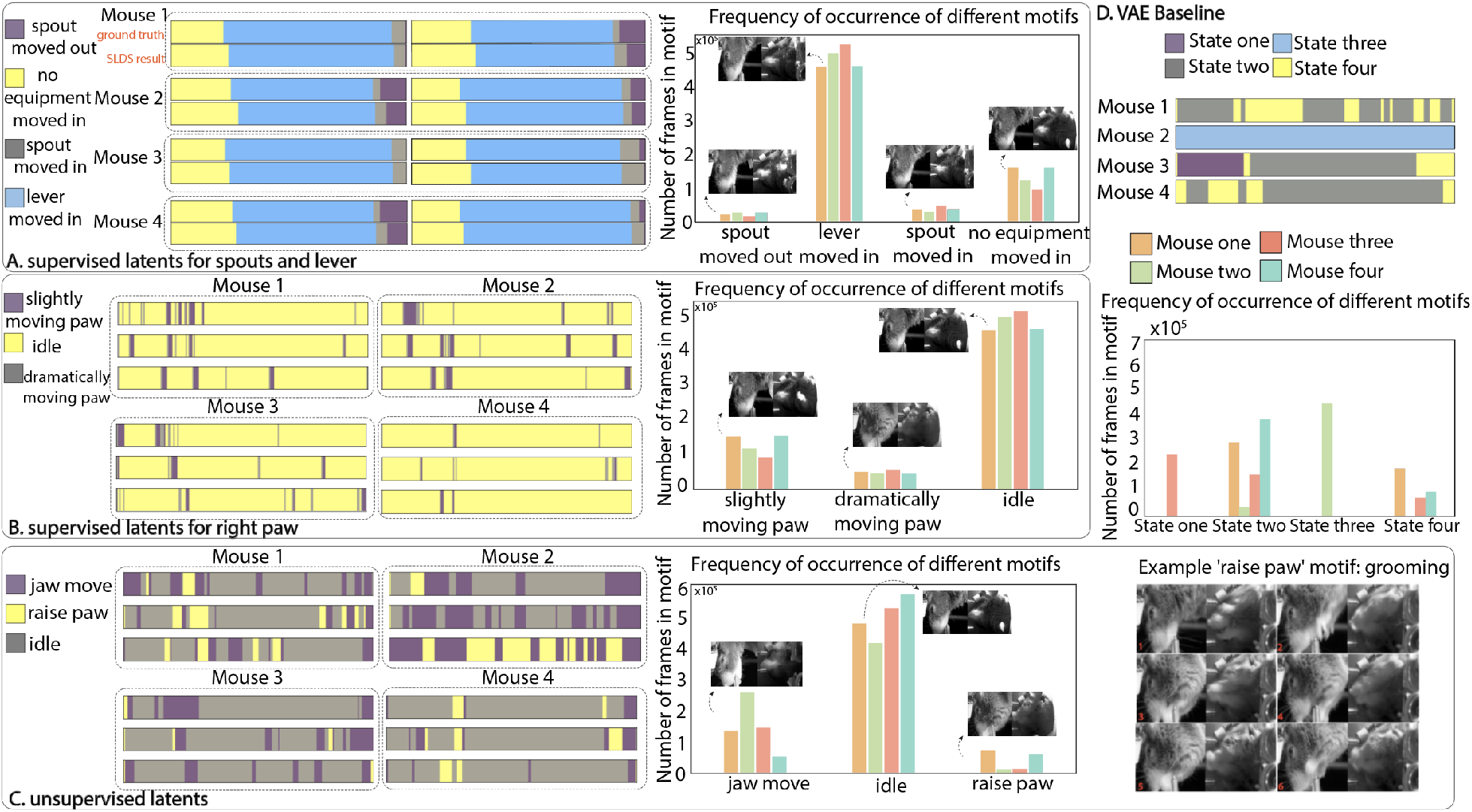
Motif generation for across-subject (supervised and unsupervised) behavioral latents using CS-VAE. SLDS results for CS-VAE latents: A. Supervised latents relating to equipment in the field of view. The equipment actions are similar for each trial. B. Supervised latents relating to tracked body parts. The ethograms for each trial across subjects and between subjects are very similar. The histogram indicates the number of frames occupied by each action per mouse. This further confirms the similarities between the supervised latents across subjects. C. Unsupervised latents also look similar across mice. Here, some example consecutive frames from the ‘raise pow’ motif are shown, which show the mouse grooming. D. As a comparison, SLDS results for the latents generated by a VAE, which failed to produce across-subject motifs.

As a baseline, we repeat this exercise on the latents of a single VAE trained to reconstruct the videos of all four mice (Fig. 5D). We see that the latents obtained by the VAE do not capture actions across subjects, and fail to cluster the same actions from different subjects into the same group.

#### Efficient Neural Decoding via Transfer Learning

To understand the relationship between neural activity and behavior, we decoded each behavioral latent with neural data across the dorsal cortex recorded using widefield calcium imaging. The decoding results for the supervised latents were similar across the CS-VAE and the PS-VAE, but we show that the neural data was also able to capture the CS-VAE unsupervised latents well (Appendix .10).

Next, as a final test of interoperability of the individual latents across mice, we used a transfer learning approach. We first trained an LSTM decoding model on 3 of the 4 mice, and then tested that model on the 4^th^ mouse while holding the LSTM weights constant but training a new dense layer leading to the LSTM (Fig. 6A, details in Appendix .10). As a baseline, we compared the performance of an individual LSTM model trained only on the 4^th^ mouse’s data. We see in Fig. 6B that, as the training set of the 4^th^ mouse becomes smaller, the transfer learning model outperforms the baseline with regards to both time and accuracy (more results and baseline comparisons in Appendix .10).

**Figure 6:**
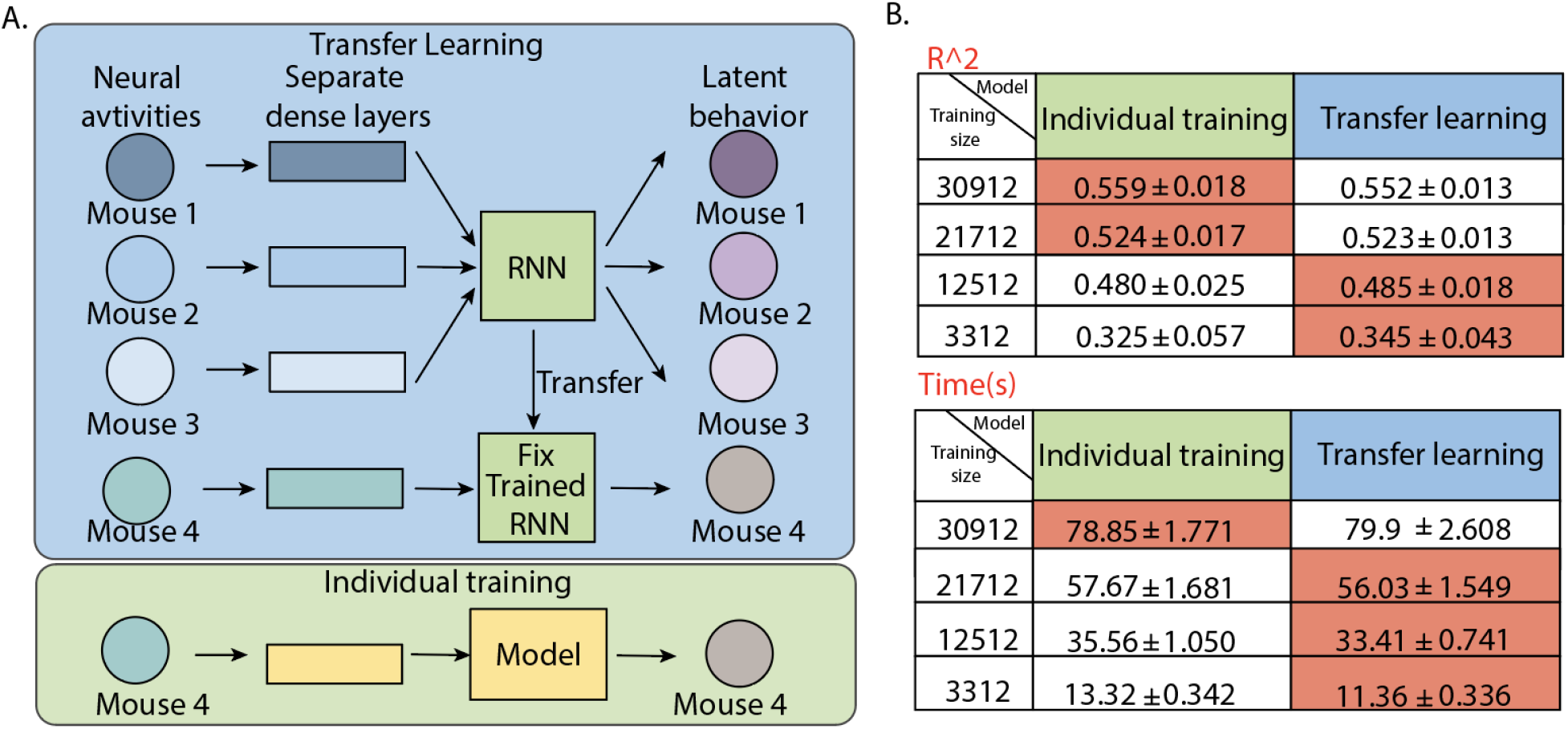
A. Transfer learning model framework. Each of the four mice has a specific dense layer for aligning the neural activities. After the model is trained using three mice, the across-subject Recurrent Neural Network (RNN) layer is fixed and transferred to the fourth mouse. As a comparison, we trained a novel RNN model for the fourth mouse and compared the accuracy with the transfer learning model B. *R*^2^ and training time trade-off for individual vs. transfer learning model as the size of the training set decreases. As the training set decreases, the transfer learning has a better performance than the individually trained model with regards to both time and *R*^2^ accuracy.

#### Neural Correlations across Mice during Spontaneous and Task-Related Behaviors

Here we explore the neural activity correlations while the subjects perform similar spontaneous behaviors vs. task-related behaviors. Across mice, we automatically identify spontaneous behaviors such as grooming and task-related behaviors such as lever pulls. We first separate the behavior from the same motif into small segments and kept the segments that have similar means and standard deviations within and across animals as shown in Fig. 7B. Next, we explore the commonalities between the neural activity of different mice as they perform these tasks by transforming the neural activity into a common subspace, using Multidimensional Canonical Correlation Analysis (MCCA). Here, we adopt the assumptions in Safaie et al. [**?**] that when the animals perform the same actions, the neural latent will share similar dynamics. We employ MCCA to align the high-dimensional neural activity across multiple subjects[**?**]. To do this, MCCA projects the datasets onto a canonical coordinate space that maximizes correlations between them (Fig. 7 C. method details in Appendix .11). Finally, we compare the commonalities across different trials in the same subject to those across subjects for different types of behaviors. In Fig. 7D, we see that for the idle behavior, the neural correlation across mice is much lower than the correlation within the same mouse; however, this does not hold for the task-related behaviors such as lever pull and licking, or the spontaneous behaviors such as grooming. For the grooming behavior, the neural correlations within and across subjects are much higher than for the idle behaviors, and in fact, even higher than the task-related behaviors. This may be due to innate behaviors having common neural information pathways across mice, whereas learnt behaviors may display significant differences across mice. Considering the region-based differences in commonalities, the sensory areas such as the visual and the somatosensory areas are much more highly correlated across mice for all behaviors as compared to motor behaviors. This may be due to the similarities in sensory feedback due to these similar behaviors but is a topic of future exploration.

**Figure 7:**
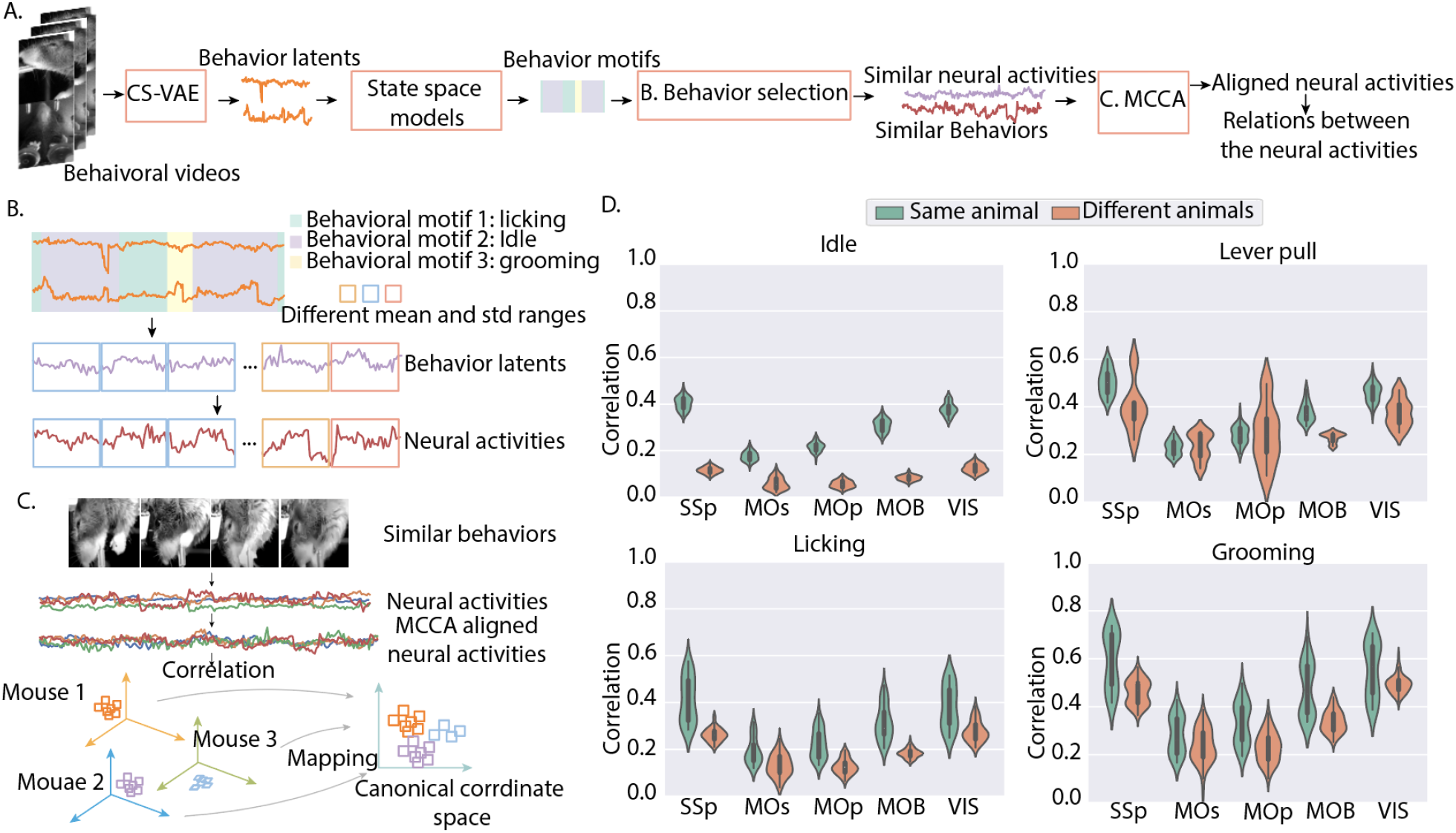
A.The overall workflow for comparing the neural activities for different subjects performing similar spontaneous behaviors: First, the behavioral videos are encoded into behavior latents by CS-VAE. Then, the behavior latents would be clustered into different motifs. After that, similar behaviors are grouped based on their mean and standard deviation values. We can therefore obtain the corresponding neural activities. Finally, the neural activities from different subjects are aligned using the MCCA. B. Behavior latents are cut into small fragments. Similar behavior fragments are grouped together based on their mean and standard deviation values. The corresponding neural activities are obtained based on the grouping results of the behavior. C. Neural activities are being aligned using MCCA. MCCA aligns the neural activities from different subjects by mapping them into the same feature spaces. D. Correlation score for behavioral-based aligned neural activity. The grooming behavior has higher neural correlation scores for cross-subjects than other behaviors.

### 2.4 Modeling Freely-Moving Social Behavior

The dataset consists of a 16 minute video of two adult novel C57BL/6J mice, a female and a male, interacting in a clean cage. Prior to the recording session the mice were briefly socially isolated for 15 minutes to increase interaction time. As preprocessing, we aligned the frame to one mouse and cropped the video (schematic in Fig. 8A; details in the Appendix .2). We tracked the nose position (*x* and *y* coordinates) of the mouse using DLC. Here, we did not include an unsupervised latent space, since the alignment and supervised labels resulted in the entire individual being explained well using the supervised latents.

**Figure 8:**
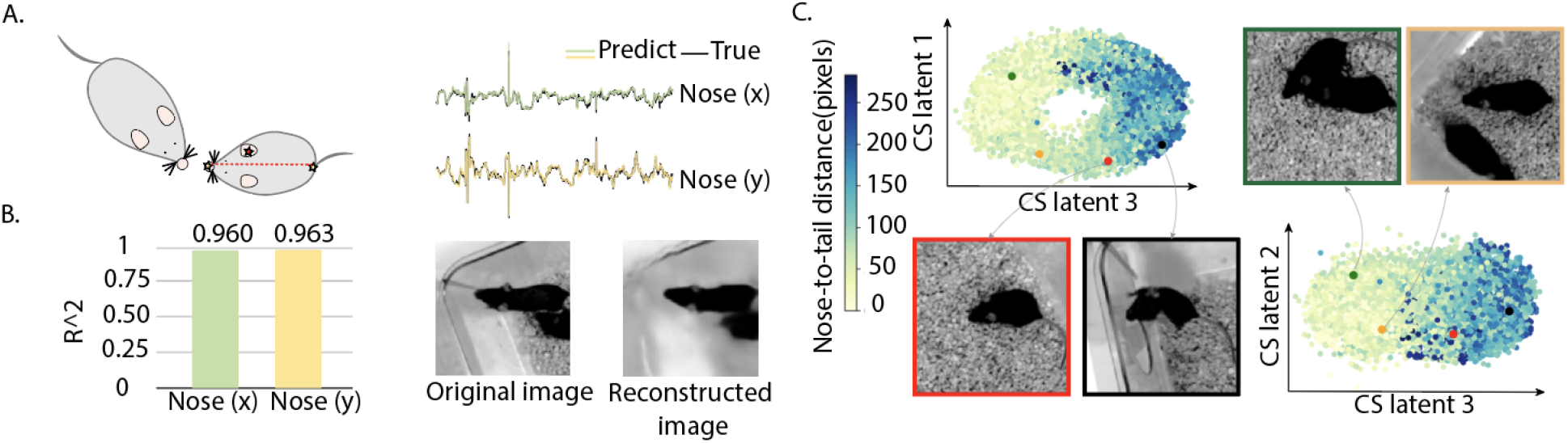
A. Image alignment for the social behavior data. B. Model performance on the social behavior dataset. C. Visualization of the CS latents overlaid with the nose-to-tail distance between the two interacting mice. The CS latents separates the frames that contain social interactions from those that do not.

#### Reconstruction Accuracy

The CS-VAE model results in a mean label reconstruction accuracy 0.961 ± 0.0017 (Fig. 8B), with the MSE for frame reconstruction as 1.21 · 10^*−*5^ (Fig. 8B). We compared the performance of our model with the VAE and PS-VAE (Table 4), and the CS-VAE model performed better than the baseline models for both image and label reconstruction. For the VAE, we obtained the *R*^2^ for nose position prediction by training a multi-layer perceptron (MLP) with a single hidden layer from the VAE latents to the nose position.

#### Disentangled Latent Space Representation

We calculated the latent traversals for each latent as in Appendix .9. As shown in the videos in Supplementary Material 4, CS latent 1 captures the second mouse to the front of the tracked mouse, CS latent 2 captures the front and above position of the second mouse, and CS latent 3 captures the position where the second mouse is below the tracked mouse.

To visualize the latent space and understand the relationship to social interactions, we plot the CS latents overlaid with the nose-to-tail distance between the two mice (nose of one mouse to the tail of the other) in Fig. 8C. We see that the CS latents represent the degree of social interaction very well, with a large separation between different social distances. Furthermore, we trained an MLP with a single hidden layer from different models’ latents to the nose-to-tail distance, and the CS-VAE produces the highest accuracy (Table 4).

#### Motif Generation

We applied a hidden Markov model (HMM) to the CS latents to uncover behavioral motifs. The three clusters cleanly divide the behaviors into social investigation vs. non-social behavior vs. non-social behavior with the aligned mice exploring the environment. To effectively visualize the changes in states, we show the ethogram in Fig. 9A. Videos related to these behavioral motifs are provided in Supplementary Material 5.

**Figure 9:**
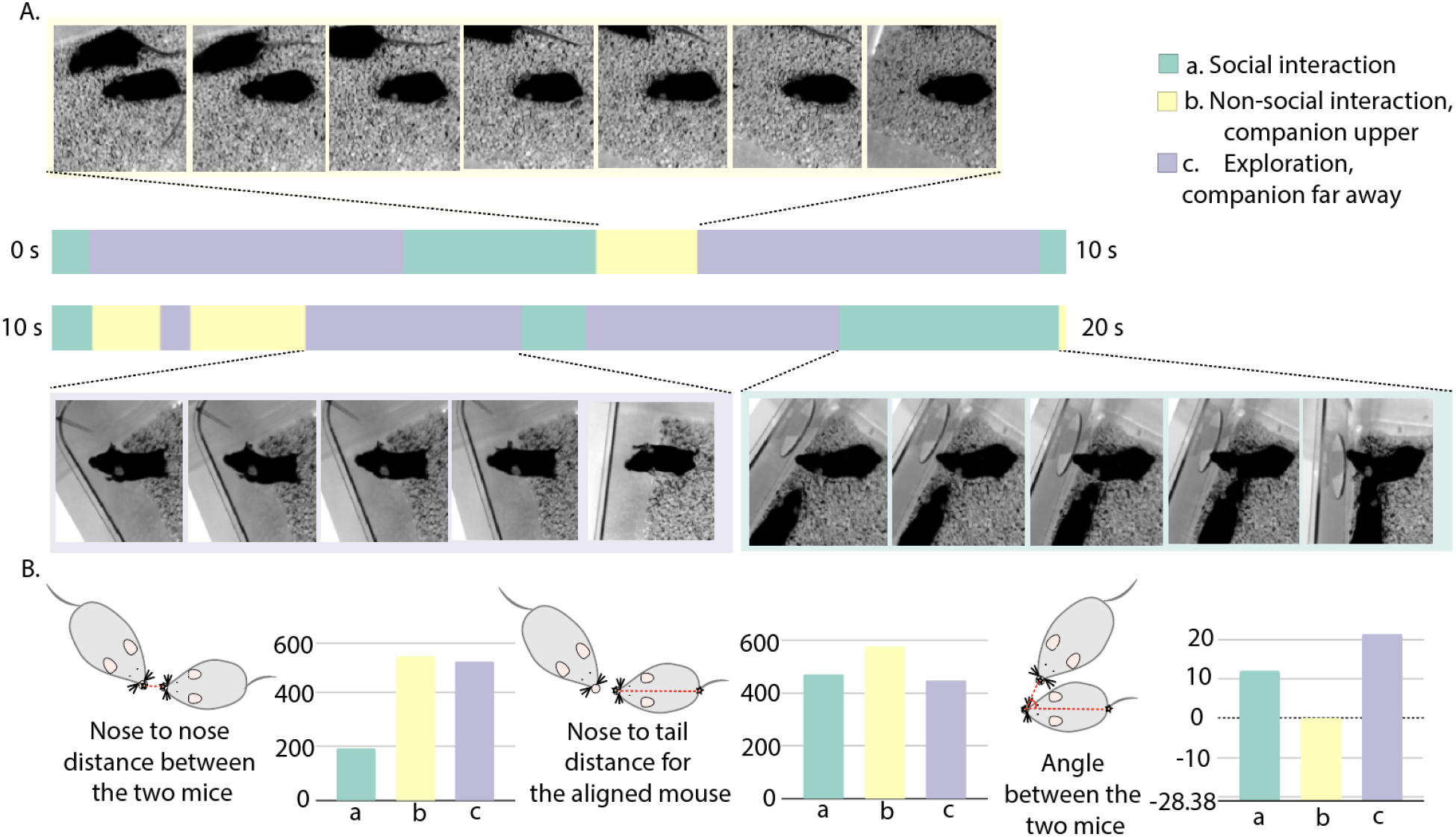
A. Ethogram for the animals’ behavior recovered using hidden Markov models (HMM) applied to the CS latents. B. Different metrics for analysing the behavioral motifs. Here, the three motifs are *a*. social interaction; *b*. non-social interaction with the companion on the upper side of the aligned mouse; *c*. non-social interaction (the aligned mouse exploring the environment with its companion far away). These metrics show the quantitative differences between the different motifs.

Lastly, we calculated different metrics to quantitatively evaluate the difference between each behavioral motif. The results are shown in Fig 9B, where we plot the average values for distances and angles between different key points. The lower distance between the two mice in *State a* demonstrates that the mice are close to each other in that state, pointing to social interactions. The smaller nose-to-tail distance for the aligned mouse in *State c* points to this state encoding for the ‘rearing’ of the mouse. The angle between the two mice further reveals the relative position between the two mice; in *State b*, the second mouse is located above the aligned mouse, while the opposite is true for *State c*. These metrics uncover the explicit differences between the different motifs that are discovered by CS-VAE.

## 3 Discussion

In the field of behavior modeling, there exist three major groups of methods, supervised, unsupervised, and semi-supervised. The supervised methods consist of methods such as DeepLabCut (DLC) [7], LEAP [6], AlphaTracker [5], amongst others. Although these methods capture the positions of the subjects, they lack the ability to model smaller movements and unlabeled behavior, and necessitate tedious labeling. On the other hand, unsupervised methods such as MoSeq [9] and Behavenet [8] lack the ability to produce intertpretable behavioral latents. While some semi-supervised methods, for instance, MSPS-VAE [2] and DBE [10], succeed in producing interpretable latents and modeling behavior across subjects, they need significant human input, and lack the ability to model freely-moving animals’ behavior. Here, we introduce a constrained generative network called CS-VAE that effectively addresses major challenges in behavioral modeling-disentangling multiple subjects and representing social behaviors.

For multi-subject behavioral modeling, the behavioral latents successfully separates the common activities across animals from the differences across animals. This behavioral generality is highlighted by the across-subject behavioral motifs generated by standard methods, and a higher accuracy while applying transfer learning for the neural decoding task. Furthermore, the SVM classification accuracy approaches 100%, which also indicates that the constrained-subspace latents well separate the differences between the subjects. In the social behavioral task, the constrained latents well capture the presence of social investigations, the environmental exploration, and the relative locations of the two individuals in the behavioral motifs. While our methods succeed in effectively modeling social behavior, it remains a challenge to separate out different kinds of social investigations in an unsupervised manner.

The constrained latents encode smoothly and discretely varying differences in behavioral videos. As seen in this work, in the across-subject scenario, the constrained latents encode the appearance of the different subjects, while in freely-moving scenario, the constrained latents capture social investigation between the subjects. The flexibility of this regularization thus gives it the ability to be fit in different conditions. Future directions include building an end-to-end structure that can captures behavioral motifs in a unsupervised way.

## 4 Methods

### Regularization of Constrained Subspace

We use the Cauchy-Schwarz divergence to regularize our constrained subspace using a chosen prior distribution. The Cauchy-Schwarz divergence *D*_*CS*_(*p*_1_, *p*_2_) between distributions *p*_1_(*x*) and *p*_2_(*x*) is given by:

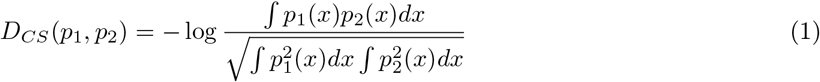

*D*_*CS*_(*p*_1_, *p*_2_) equals zero if and only if the two distributions *p*_1_(*x*) and *p*_2_(*x*) are the same. By applying the Parzen window estimation technique to *p*_1_(*x*) and *p*_2_(*x*), we get the entropy form of the Equation [11]:

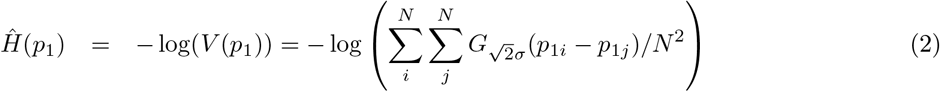

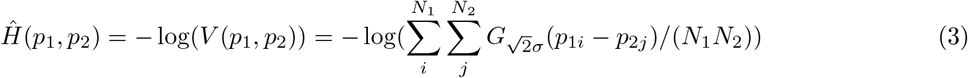

Here, *p*_1*i*_ represents the *i*th sample from the distribution *p*_1_, i.e., *p*_1_(*x*_*i*_). − log(*V* (*p*_1_)) and − log(*V* (*p*_2_)) are the estimated quadratic entropy of *p*_1_(*x*) and *p*_2_(*x*), respectively, while − log(*V* (*p*_1_, *p*_2_)) is the estimated cross-entropy of *p*_1_(*x*) and *p*_2_(*x*). *G* is the kernel applied to the input distribution; here it is chosen to be Gaussian. *N, N*_1_, and *N*_2_ are the number of samples being input into the model while *σ* is the kernel size. The choice of the kernel size depends on the dataset itself; generally, the kernel size should be greater than the number of the groups in the data. Equation (1) can be expressed as:

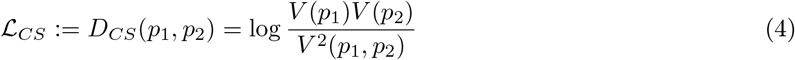

Here, *p*_1_(*x*) represents the distribution of our CS latent space, and *p*_2_(*x*) the chosen prior distribution. In Equation (4), minimizing *V* (*p*_1_) would result in the spreading out of *p*_1_(*x*), while maximizing *V* (*p*_1_, *p*_2_) would make the samples in both distributions closer together [11]. Thus, we minimize this term in the objective function while training the model. However, it may be necessary to stop at an appropriate value, since overly spreading out *p*_1_(*x*) may lead to the separation of the samples from the same groups, while making *p*_1_ and *p*_2_ excessively close may cause mixtures of data points across groups.

In short, the Cauchy-Schwarz divergence measures the distance between *p*_1_ and *p*_2_. In our work, we adopt a variety of distributions as a prior distribution *p*_2_(*x*), and we aim to project the constrained subspace latents onto the prior distribution (see Fig. 1).

### Optimization

The loss for the CS-VAE derives from that for the PS-VAE, and is given by:

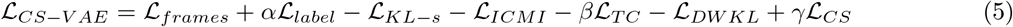

Here, the terms ℒ_*frames*_ and ℒ_*label*_ represent the reconstruction loss of the frames and the labels, respectively. The ℒ_*KL−s*_ represents the KL-divergence loss for the supervised latents while ℒ_*ICMI*_, ℒ_*T C*_, and ℒ_*DW KL*_ form the decomposed version of the KL loss for the unsupervised latents. Lastly, the ℒ_*CS*_ represents the CS-divergence loss on our constrained latents. *α* is introduced to control the reconstruction quality of the labels, *β* is adopted to assist the model in producing independent unsupervised latents, and *γ* is implemented to control the variability in the constrained latent space for better separation. The detailed explanations and derivations for each term in the objective function are in Appendix .3. Furthermore, the loss terms in Equation (5) can be modified to fit various conditions. For a freely-behaving social task, the background for one individual in the container could be the edge of the container as well as the rest of the individuals in the container. The choice of hyperparameters and the loss curves through the training process is shown in Appendix .5 and .7, respectively.

### Visualization of the latent space

To test how the image varies with a change in the latent, one frame from the trials is randomly chosen as the ‘base image’, and the effect of varying a specific latent at a time is visualized and quantified. This is known as the ‘latent traversal’ [2]. First, for each latent variable, we find out the maximum value that it occupies across a set of randomly selected trials. We then change that specific latent to achieve its maximum value, and this new set of latents forms the input to the decoder. We obtain the corresponding output from the decoder as the ‘latent traversal’ image. Finally, we visualize the difference between the ‘latent traversal’ image and the base image. The above steps are performed for each latent individually. In videos containing latent traversals (Supplementary Material), we change the latent’s value from its minimum to its maximum across all trials, and input all the corresponding set of latents into the decoder to produce a video.

### Behavioral Motif Generation

Clustering methods such as Hidden Markov Models (HMM) and switching linear dynamical systems (SLDS) have been applied in the past to split complex behavioral data into simpler discrete segments [16] (see Appendix .6 for details). We use these approaches to analyze motifs from our latent space, and directly input the latent variables into these models. In the case of multi-subject datasets, our goal is to capture the variance in behavior in a common way in the across-subject latents, i.e., recover the same behavioral motifs in subjects performing the same task. In the case of freely-moving behavior, our goal is to capture motifs related to social behavior.

### Efficient Neural Decoding

Decoding neural activity to predict behavior is very useful in the understanding of brain-behavior relationships, as well as in brain-machine interface tasks. However, models to predict high-dimensional behavior using large-scale neural activity can be computationally expensive, and require a large amount of data to fit. In a task with multiple subjects, we can utilize the similarities in brain-behavior relationships to efficiently train models on novel subjects using concepts in transfer learning. Here, we represent across-subject behavior in a unified manner and train an across-subject neural decoder. Armed with this across-subject decoder, we show the decoding power on a novel subject with varying amounts of available data, such that it can be used in a low-data regime. The implementational details for this transfer learning approach can be found in Appendix .10.

### Behavior election for innate behaviors studying

While the behavioral features extracted from the previous sections are successful in capturing similar spontaneous behaviors across various animals, the behavioral patterns within the same motifs can exhibit substantial variation. For instance, in the case of the raising paw motif, continuous movement of the paws could be indicative of either grooming or other complex behaviors. To overcome this challenge, we divided the behaviors belonging to the same motif into smaller segments and calculated the corresponding mean and standard deviation of the behavioral latents. Subsequently, we compared these values and retained the segments that exhibited similar mean and standard deviations both within and across animals, as illustrated in Fig. 7B. These steps were repeated for all the behavioral motifs examined in this study.

In addition to the spontaneous behaviors discussed above, we also selected an ‘idle’ behavior that captured the mouse’s inactivity and a task-related behavior, namely the ‘lever pull’ behavior, which signaled the initiation of each task.

## 5 Appendix

### .1 Experimental Methods and Preprocessing for the Multi-Subject Dataset

In our work, we employed a subset of the behavioral dataset detailed in Musall et al., 2019 [14]. Briefly, the task entailed pressing a lever to initiate the task, after which a visual stimulus was displayed towards the left or the right. After a delay period, the spouts come forward, at which time the mouse makes its decision by licking the spout corresponding to the direction of the visual stimulus (left or right). Finally, the mice receive a juice reward if they choose correctly.

We tested the CS-VAE on the behavioral data for the four mice performing a visual task and randomly chose 388 trials per mouse each of the trials has a 189 number of frames. Each frame was pre-processed and resized to have both the length and width being 128. One example trial for each mouse can be found in Supplementary Material 1.

Before inputting the data into the model, we sorted the trials by the amount of variance in the images, and shuffled the first half (high variance) and the second half (low variance) of the dataset separately. This was done to speed up training by training the model on high-variance trials first. We tested our model by randomly choosing 4 trials from all trials for each mouse 5 times. The same procedure was applied when training the model on the simulation dataset, i.e., the doctored data for one subject.

### .2 Experimental Methods and Preprocessing for the Freely-Moving Social Behavior Dataset

The dataset consists of a 16-minute video of two adult novel C57BL/6J mice, a female and a male, interacting in a clean cage. Prior to the recording session, the mice were briefly socially isolated for 15 minutes to increase interaction time. This dataset was collected by one of the authors. The original data has 24917 number of frames with length and width being 1920 and 1080, respectively. The example fraction of the video can be found in Supplementary Material 6.

The nose, ears, and tail base of each mouse were manually annotated using AlphaTracker. We kept 19659 number of frames that have the labels for preprocessing and training. We perform several preprocessing steps to align and crop the video as well as the labels based on one of the two mice (Mouse 1, female). All of the preprocessing steps were based on the AlphaTracker labels. For each frame, we first rotate it to ensure that the nose and tailbase for Mouse 1 are on the same horizontal line, with the central point for rotation as the left ear. Next, we aligned the frame such that the left ear of Mouse 1 was at the same location across all frames. Finally, we resize the frame to be 128 × 128 and consequently the AlphaTracker labels. For this dataset, since there was a relatively low number of frames, we obtained the CS-VAE MSE and label *R*^2^ for the entire dataset.

### .3 Methodological details of the Partitioned Subspace VAE

The Partitioned Subspace VAE (PS-VAE) was introduced in [2], and we borrow the notation used in that paper when detailing the CS-VAE. Thus, we include here a full description of the model.

First of all, we define the input frame as *x*, and the corresponding pose estimation tracking label as *y*. The reconstructed variables are termed 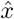 and *ŷ*, respectively. The supervised latent space is denoted as *z*_*s*_, unsupervised latent as *z*_*u*_, and the background latent as *z*_*b*_. In a VAE model, we would like to minimize the distance, typically the KL divergence, between the posterior distribution of the latent variables *p*(*z*|*x*) and a chosen distribution *q*(*z*|*x*). However, since *p*(*z*|*x*) is an unknown distribution, the Evidence Lower Bound (ELBO) is introduced as an alternative method to reduce the KL divergence:

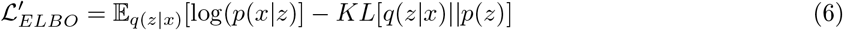

Following [2], if we have a finite dataset 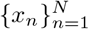, and we treat *n* as a random variable with a uniform distribution *p*(*n*) while defining *q*(*z*|*n*) := *q*(*z*_*n*_|*x*_*n*_), we can rewrite the ELBO as:

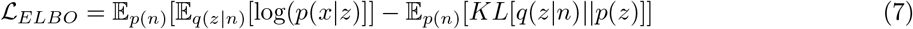

We define the loss over frames ℒ_*frames*_ as the first of the two terms above. In the PS-VAE model, there are two inputs: frames *x* and labels *y*. Therefore, in Equation (7), instead of writing the input likelihood as *p*(*x*|*z*), we can now write it as *p*(*x, y*|*z*). A simplifying assumption is made that *x* and *y* are conditionally independent given *z*, and thus we can directly write ℒ_*frames*+*labels*_ as ℒ_*frames*_ + ℒ_*labels*_, where ℒ_*labels*_ is calculated by replacing *x* with *y* in ℒ_*frames*_.

After assuming the prior *p*(*z*) has a factorized form: *p*(*z*) = Π_*i*_ *p*(*z*_*i*_), the KL term ℒ_*KL*_ can be split as the addition of *ℓ*_*KL−s*_ and *ℓ*_*KL−u*_, i.e., the KL terms for the supervised and unsupervised latents, respectively. We decompose the KL term for the unsupervised latent as the following [2].

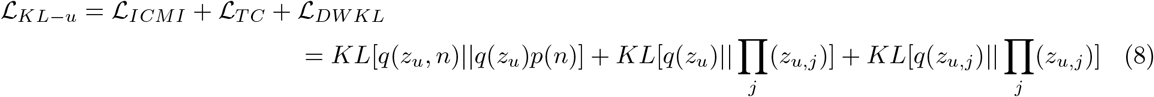

where *j* represents the latent dimension, ℒ_*ICMI*_ is the index-code mutual information, which measures how well the latent encodes the corresponding input data. The term TC is short for total correlation, which measures the interdependency of each latent dimension. The third term, ℒ_*DW KL*_ is the dimension-wise KL, which calculates the KL divergence for each dimension individually. Finally, the resulting subspace is forced to be orthogonal by applying orthogonal weights across all the different latents.

The authors in [2] introduce an extension to PS-VAE for modeling multi-session data. The Multi-Session PS-VAE (MS-PS-VAE) can only work with a labeled set of discrete sessions, as described in the Introduction. The images from each session are labeled, and the session-specific latents are enforced to be static over time, thus capturing the image-related details. To enforce the background latents to be static over time in a particular session, and to maximize the difference in the background latents across different sessions, the triplet loss is introduced in MS-PS-VAE. As described in the Introduction, this loss term artificially places the latents from the same session together while separating the latents from different sessions. The triplet loss is computed as the following.

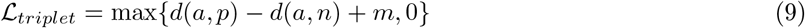

Here, *a* is the anchor point, *p* is the positive point, *n* is the negative point, and *m* is a margin. The function pulls the point *p* towards point *a*, and pushes the point *n* away from point *a*. While training, the data from multiple sessions is included in each mini-batch. The data from each session is split in three, and each third from the same session acts as an anchor and positive point, while the data from another session acts as a negative point. Practically, this requires as many sessions as possible in the same mini-batch during the training for accurate results. As the number of sessions increases, this method becomes computationally intractable, and may lead to unsatisfactory reconstruction results. Moreover, this loss does not allow for varying backgrounds across any one session.

In the MS-PS-VAE model, the triplet loss was applied as a supervised manner to pull the data from the same subject being closer while pushing the different subjects away from each other. This method is only useful when the number of sessions is known, and is not applicable in an open-field setting, for example while modeling freely-moving social behavior as in this manuscript.

Therefore, in this manuscript, we introduce a regularization term that can automatically separate different subjects in the background latent space without specifying the number of sessions or labeling each frame as belonging to a specific session.

### .4 Model Architecture and Training

Our computational experiments were carried out using TensorFlow and Keras. The image decoder we use is symmetric to the encoder, with both of them containing 14 convolution layers. We applied the Adam optimizer with learning rate as 10^*−*4^. For the multi-subject dataset, we fixed our batch size to be 256 and trained for 50 epochs. For the freely-moving social behavior dataset, we trained for 500 epochs with batch size 128.

### .5 Choice of Hyperparameters

In the multi-subject dataset, four coefficients need to be decided for the objective function as indicated above: {*α, β, σ, γ*}. There is a balance between the choice of *β* and *γ*: properly choosing the values could separate the latent in the unsupervised space and the latents in both unsupervised and background space as well. A large separation of the background latent may potentially lead to unsatisfactory reconstruction results. The choice of kernel size *σ* depends on the dataset, and should be larger than the number of distinct groups in our dataset; since in our current experiments, we have at most four groups, we set *σ* = 15. Moreover, we set *α* to 1000, *β* to 5, *γ* to 500. We set the dimensionality of the supervised latent space equal to the number of tracked video parts, which is 5 in our case. We set the dimensionality of the unsupervised latent space as 2, while that of the background latent space as 2.

In the social behavior task, we track the nose location as the supervised latent, since the other labels do not have a high variance (due to the alignment process). Additionally, we do not need any unsupervised latents to explain the individual’s behavior. The CS latent in this setting has 3 dimensions. Here, *α* is 1200, *γ* is 200, and the kernel size is 20.

The hyperparameters chosen for all three datasets are shown in Tables 2 and 3.

**Table 1:**
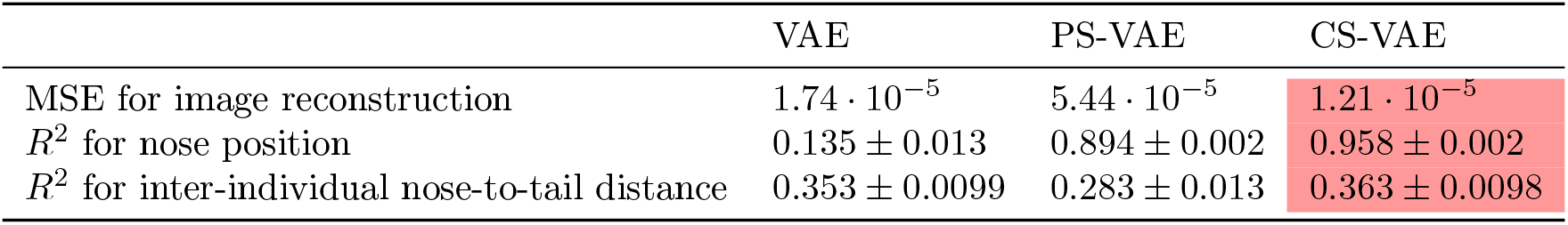
Comparison of different models on the freely-moving social behavior dataset

|  | VAE | PS-VAE | CS-VAE |
| --- | --- | --- | --- |
| MSE for image reconstruction | $1.74 \cdot 10^{-5}$ | $5.44 \cdot 10^{-5}$ | $1.21 \cdot 10^{-5}$ |
| $R^2$ for nose position | $0.135 \pm 0.013$ | $0.894 \pm 0.002$ | $0.958 \pm 0.002$ |
| $R^2$ for inter-individual nose-to-tail distance | $0.353 \pm 0.0099$ | $0.283 \pm 0.013$ | $0.363 \pm 0.0098$ |

**Table 2:**
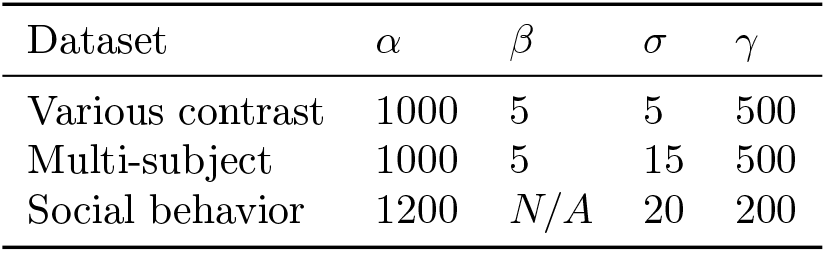
Hyperparameter for different dataset

**Table 3:**
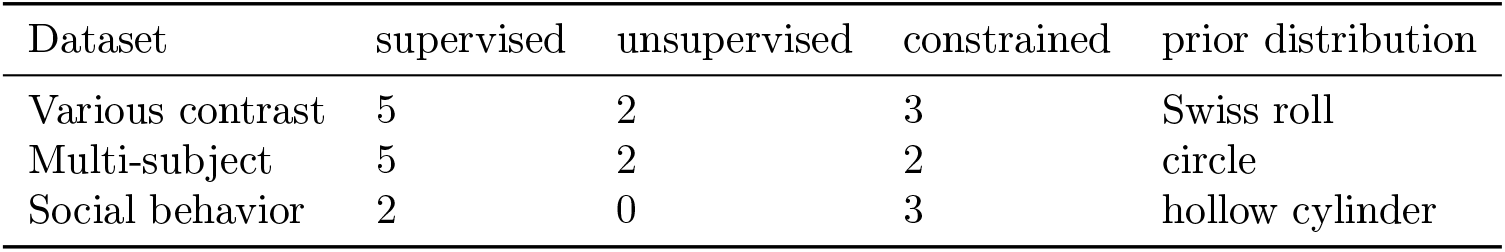
Latent dimensions and the prior distribution for different dataset

**Table 4:**
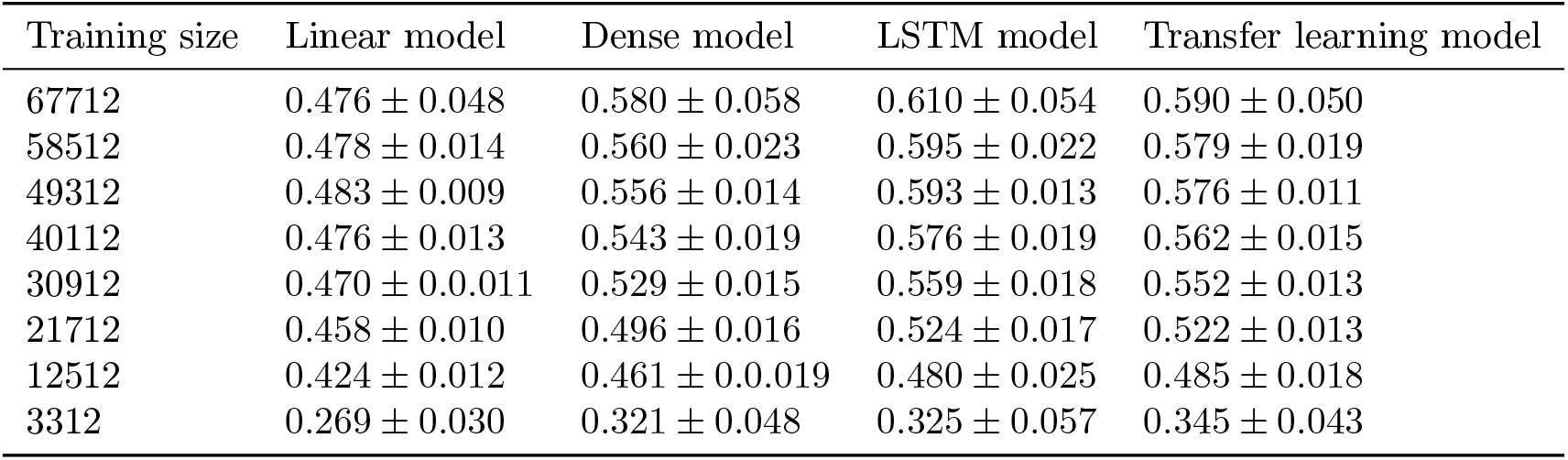
Training size vs *R*^2^ value for multi-subject dataset

### .6 Motif Generation

A switching linear dynamical system (SLDS) consists of discrete latent state *z*_*t*_ ∈ {1, 2, ..*K*}, continuous latent state *x*_*t*_ ∈ R^*M*^, and the observation state *y*_*t*_ ∈ R^*N*^. Here, *t* = 1, 2, 3, .., *T* is the time step, *T* is the length of the input signal; *K* is the number of discrete states; *M* is the number of latent dimensions; *N* is the observation dimensions. The discrete latent state *z*_*t*_ follows the Markovian dynamics with the state transition matrix expressed as:

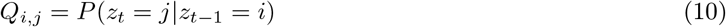

The continuous latent state *x*_*t*_ has the following linear dynamical relations that determined by *z*_*t*_.

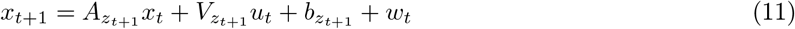

Here, 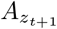 is the dynamic matrix at state *z*_*t*+1_; *u*_*t*_ is the input at time t, with 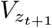 being the control matrix; 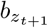 is the offset vector and *w*_*t*_ being the noise which is generally the zero mean Gaussian. Here, our observation model is in Gaussian case; therefore, the observation *y*_*t*_ is expressed as:

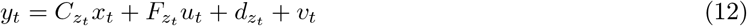

Here, 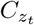 is the measurement matrix at state *z*_*t*_; 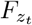 is the feedthrough matrix which directly feed the input into the observation; 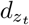 is the offset vector and *v*_*t*_ is the noise. Here the update was accomplished by the Expectation-Maximization(EM) algorithm. In the E-step, the model updates the hyperparameters. In the M-step, the log-likelihood in Eq.12 is being maximized.

To implement the SLDS, we adopted the open source software from Linderman et al.[16]. We fit the SLDS using different latent dimensions, where the observation dimension was the order of latent dimension and the number of states was determined by visualizing the videos. We use SLDS’s to model the motifs in the multi-subject dataset since the behaviors are well separated using their dynamics. We use K-means to model the motifs in the freely-moving social behavior dataset since the behaviors are well separated directly in state space. An autoregressive HMM (a simpler model than an SLDS) applied to the CS latents in the social behavior dataset leads to similar results as the K-means.

### .7 Loss Curves

We show the learning curve for each loss term for both dataset to precisely quantify the model, in Fig. 10. For the multi-sujbect dataset (Fig. 10A), for the unsupervised latents, the final loss for dimension-wise KL, total correlation, and the mutual information are 11.7, −4.8, and −4.6, respectively. The final KL loss for the supervised latents is 5.06 and the final CSD loss for the CS latents is 0.1. For the free behaving dataset, the loss curves for each loss term are shown in Fig. 10B. By the end of the training process, the KL loss for the supervised latents is 7.01 and the CSD loss for the CS latents is 1.15.

**Figure 10:**
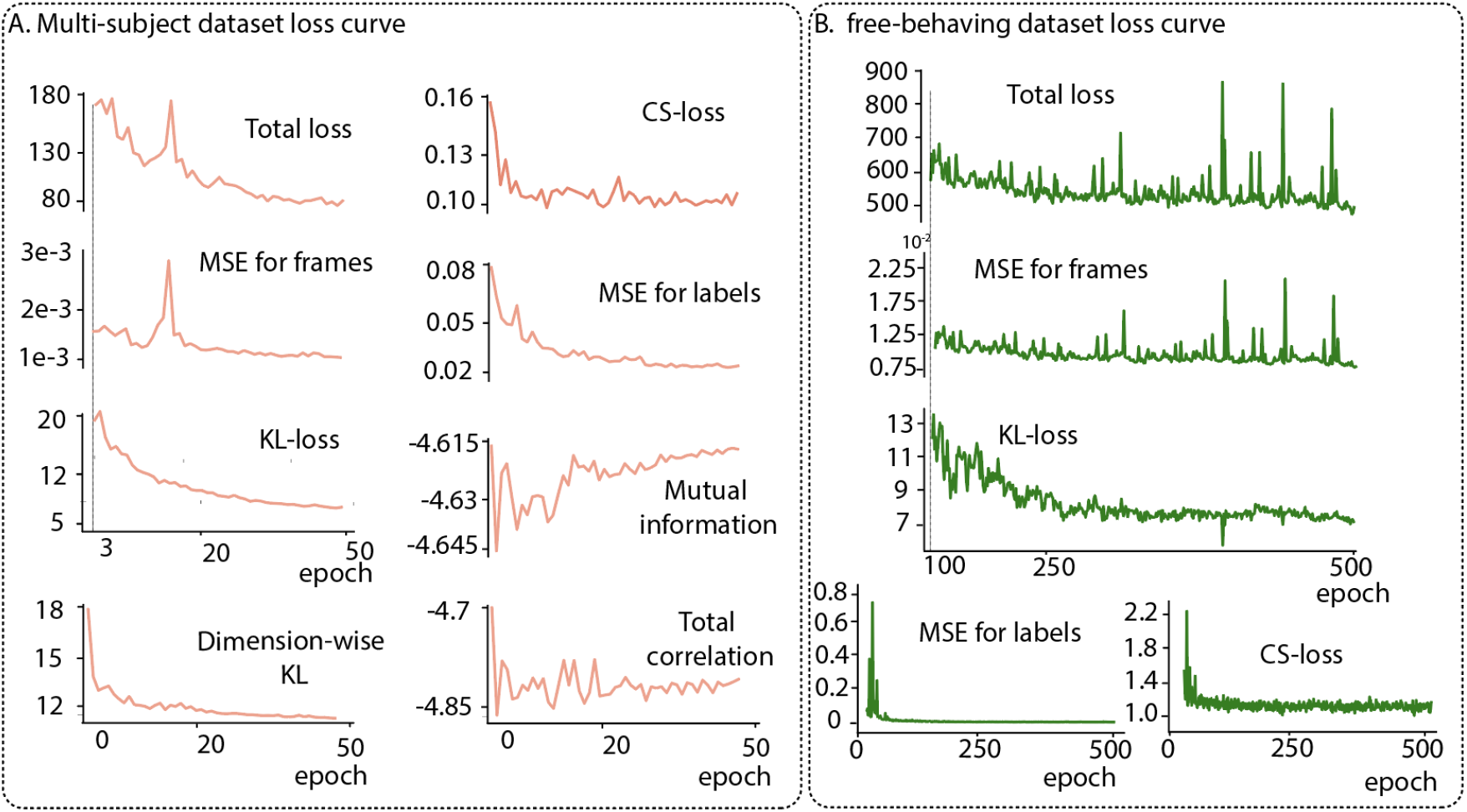
Loss curve for A. training the multi-subject dataset B. training the freely behaving dataset with the specified hyperparameters as in Tables 1 and 2.

### .8 SVM

To further quantify the separation of the latents between different subjects, we applied a supervised classification method to decode the identity of the subject using each latent.

After randomly shuffling all the latents, we split all the trials into training trials and test trials, with each mouse having 368 trials in the training set and 20 trials in the test set, and repeated this 5 times with different random seeds.

### 9 Latent traversal

For the multi-subject dataset, we tested the latent traversal with the same base image to validate the results, shown in Figure 11. Here, we randomly chose a frame from a mouse and changed each individual latent within different ranges as detailed in the Methods. For example, in Figure 11, the first row contains the output when the corresponding latent is changed to take on the maximum value from the range of Mouse 1. Similar to the figures in the main text, the upper images are the latent traversal images while the lower ones are the difference between the upper and original images. We see that the base image from Mouse 3 can be flexibly changed to produce a different mouse when changing the CS latent. Moreover, when changing the supervised and unsupervised latents for the different mice, Mouse 3 seems to be flexibly changing with these latents from different mice.

**Figure 11:**
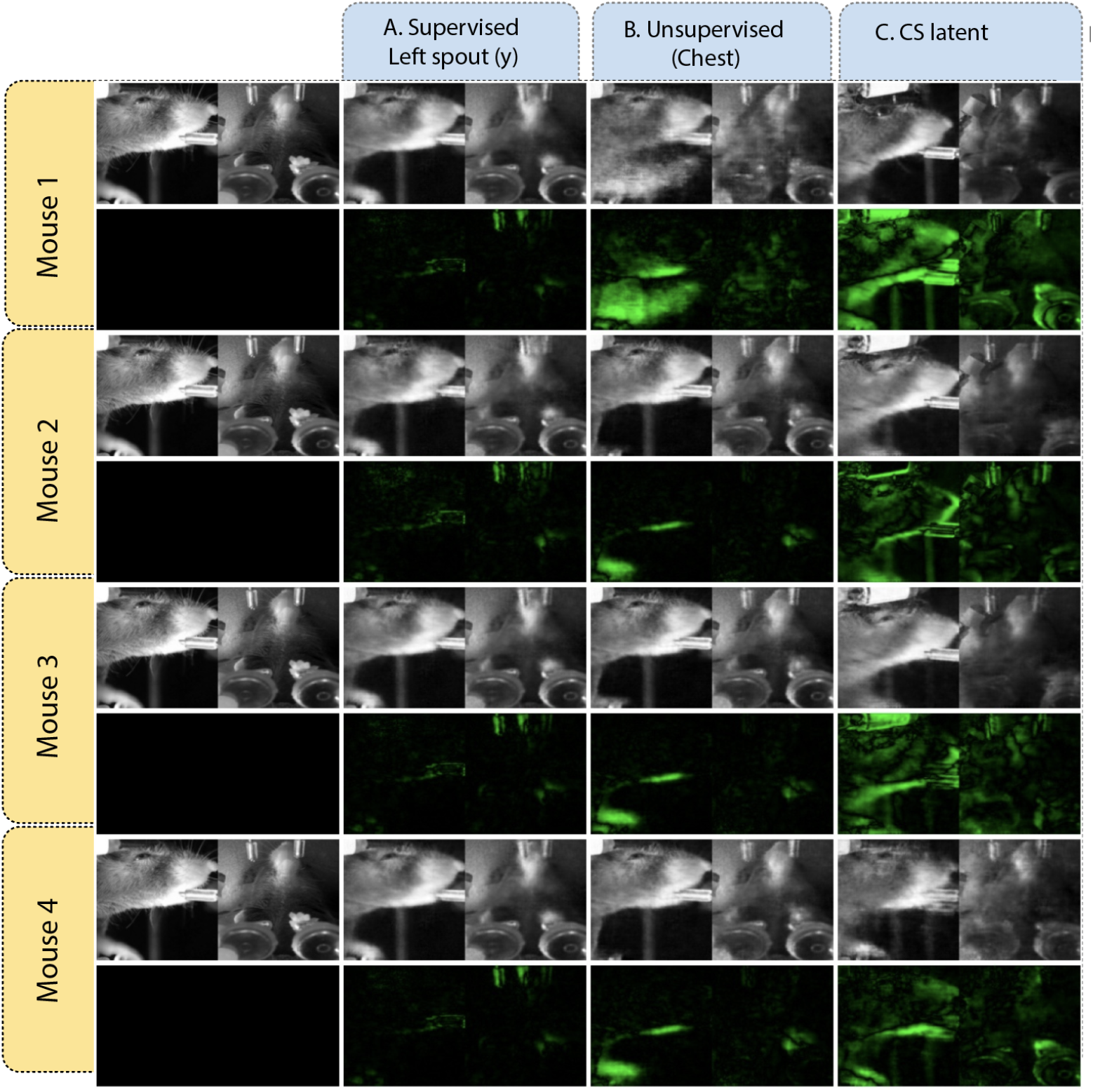
Latent traversals for the multi-subject dataset for the four mice with the same base image A. an example supervised latent, B. an example unsupervised latent, and C. an example CS latent. We see that the same base image (Mouse 3) is transformed into a different mouse each time when changing the CS latent.

To better visualize the specialization of each latent, we generated the latent traversal videos for each latent with different base images. For different mice, we, first of all, find the maximum and the minimum value for the specific latent. Then, change the latent within that range with 0.5 per step. Finally, concatenate all the latent traversal images into videos. The videos can be found in Supplementary Material 3.

We performed a similar visualization on the freely-moving social behavior dataset for the CS latents. The latent traversal videos can be found in Supplementary Material 4, and some clips from the videos are shown in Fig 12.

**Figure 12:**
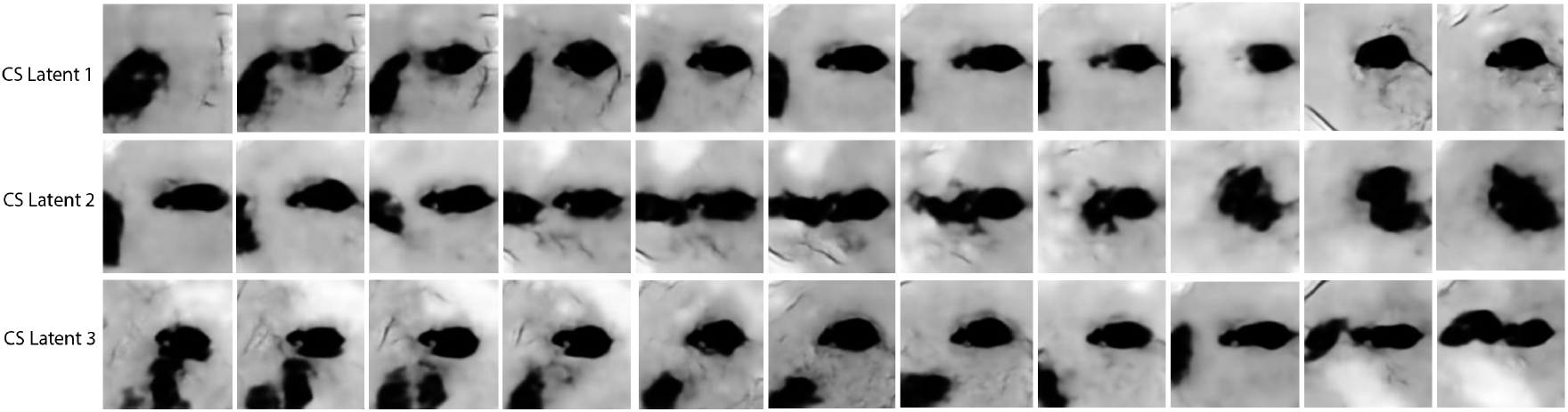
Latent traversals on the CS-latents for the freely-moving social behavior dataset. We see that the latents all encode for social interactions between the two mice.

**Figure 13:**
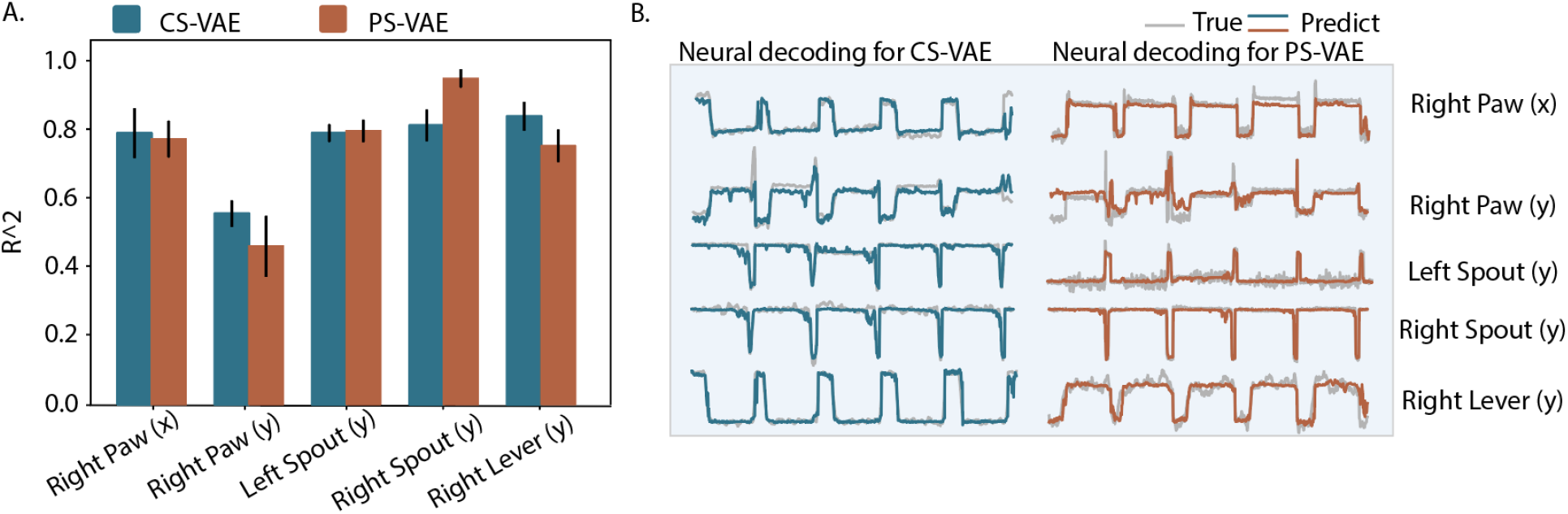
Neural decoding for CS-VAE vs. PS-VAE.

### .10 Neural decoding models

The trials were first shuffled and then split into training and testing. Next, we employed the CS-VAE generated latent representations, and choose one example subject to decode the behavior at time *t* using the neural activity recorded between *t* − 0.15s and *t*. We applied four types of models to compare the performance. A linear model which directly maps the neural activities into the behavior. A multilayer perceptron (MLP) with three dense layers to train the decoder. We used the Adam optimizer with learning rate decay from 0.1 with 0.3 decay rate for every 5 step. The batch size was fixed to be 150 and trained for 200 epochs. A LSTM model, which begin with a dense layer followed by a LSTM layer with a drop-out rate being 0.5 and another dense layer at the end. We applied the same training strategy as in MLP model.

We introduced a model based on transfer learning to perform the decoding test on the previously tested subject. The rest of the three mice were the input to the original training model. The procedures were similar to before, after the trials were shuffled and split, we decoded the behavior directly with the raw neural activities with the time window being 0.15*s*. After that, we implemented three perceptron layers for each of the three mice before the output of which went into a recurrent neural network (RNN). The RNN consisted of one long short-term memory (LSTM) layer with a unit number of 64 and a drop-out layer with a rate being 0.5. We applied the Adam optimizer with learning rate decay from 0.1 with 0.3 decay rate for every 5 step. The batch size was 150 and we trained for 200 epochs. After we finished training the original network, we transferred the RNN model to the new model which was applied to train the fourth mouse alone. For the fourth mouse, the trials were split with different training and testing ratios. After applying the same steps to the data, the neural activities then went through a new perceptron layer before going through the pre-trained RNN model. We applied the Adam optimizer with the same learning rate decay procedures as well. We again, trained for 200 epochs with batch size being 128 this time. The trade-off between accuracy and time for different models can be found in Tables 4 and 5.

**Table 5:**
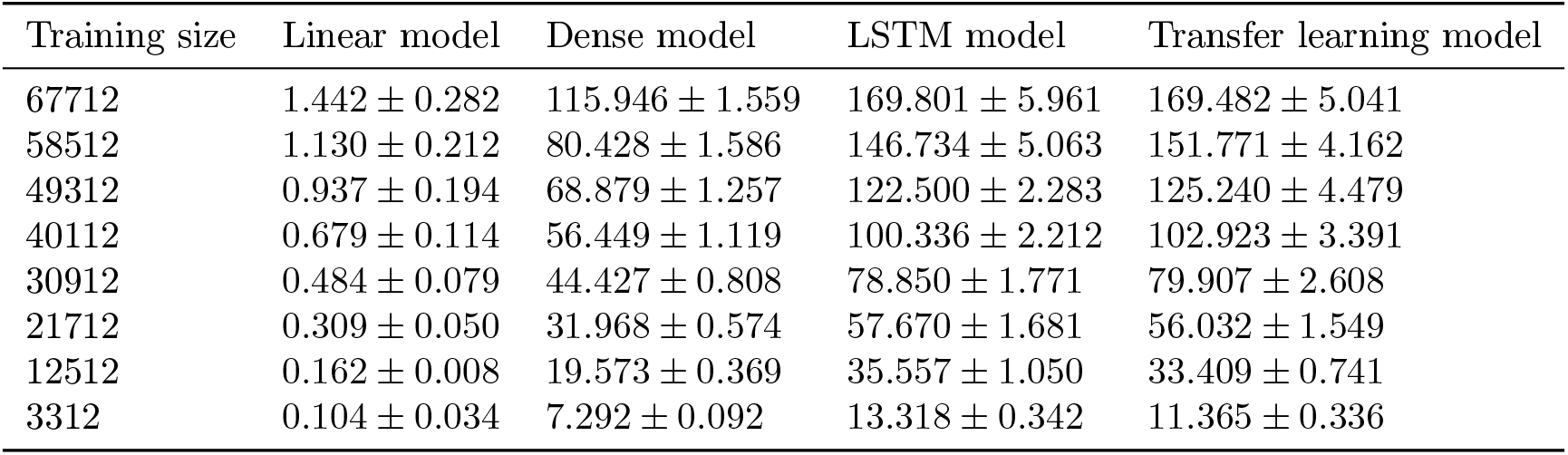
Training size vs time usage for multi-subject dataset

### .11 Multidimensional Canonical Correlation Analysis (MCCA) for neural signal alignment

In our work, after extracting similar behaviors chunks from different individuals, we then extracted the corresponding neural activity for each subject. To smooth away the discreteness of the neural activity chunks, we shuffled the chunks before concatenating them together. After that, we performed the MCCA for all four subjects on each brain region. For each brain region, we choose the four sets of neural activities being the same length *d, X*1 = {*x*1_1_, *x*1_2_, …, *x*1_*n*_} ∈ *R*^*n×d*^, *X*2 = {*x*2_1_, *x*2_2_, …, *x*2_*n*_} *R*^*m×d*^, *X*3 = {*x*3_1_, *x*3_2_, …, *x*3_*n*_} ∈ *R*^*k×d*^, and *X*1 = {*x*1_1_, *x*1_2_, …, *x*1_*n*_} *R*^*l×d*^. Here, we choose the minimum number of region dimensionality in all of the four subjects as the dimension of canonical coordinate space, *minimum* {*n, m, k, l*}, and is annotated as *j*. For each dimension, define the projection weights for each dataset as *a*_*j*_ = {*a*_*j*1_, *a*_*j*2_, .., *a*_*jn*_ }, *b*_*j*_ = {*b*_*j*1_, *b*_*j*2_, .., *b*_*jn*_}, *c*_*j*_ = {*c*_*j*1_, *c*_*j*2_, .., *c*_*jn*_}, and *d*_*j*_ = {*d*_*j*1_, *d*_*j*2_, .., *d*_*jn*_}. The resulting projected datasets are now *d*-dimensional arrays: *u*1_*j*_ = ⟨*a*_*j*_, *X*1⟩, *u*2_*j*_ = ⟨*b*_*j*_, *X*2⟩, *u*3_*j*_ = ⟨*c*_*j*_, *X*3 ⟩, and *u*4_*j*_ = ⟨*d*_*j*_, *X*4⟩. For each of the coordinate spaces, the objective functions can be written as:

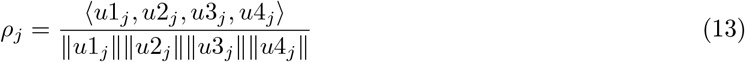

Generally, for each pair of canonical components, the above equation is solved iteratively to find the best projects that can maximize the correlation. During training, the orthogonality between each canonical component is constrained. In our experiment, we calculated the across-subject correlations for each obtained CCs and kept the highest correlation value for each pair, here termed *ρ*_1_ (Equation 13). We performed the above task for each brain region. In addition, we shuffled the chunks ten times and repeated the above steps. We also calculated the canonical component for the same subject having similar behaviors. We applied the same methods as stated above to find similar behavior components and the corresponding neural activities. We divided the obtained neural activities into two parts with the same length and performed the CCA on those two signals. We calculated the correlation between the first two canonical correlation axes as the baseline.

### .12 Code

The code for training the CS-VAE can be found in Supplementary Material 7. The code can be executed by simply compiling the script ‘train.py’. All the code are available at: https://github.com/saxenalabneuro/Behaivoral-feature-extraction-CS-VAE.

