## Supplementary Info for "Disentangled multi-subject and social behavioral representations through a constrained subspace variational autoencoder (CS-VAE)"

$$\mathcal{L}'_{ELBO} = \mathbb{E}_{q(z|x)}[\log(p(x|z)] - KL[q(z|x)||p(z)] \tag{1}$$

Following [2], if we have a finite dataset $\{x_n\}_{n=1}^{N}$, and we treat $n$ as a random variable with a uniform distribution $p(n)$ while defining $q(z|n) := q(z_n|x_n)$, we can rewrite the ELBO as:

$$\mathcal{L}_{ELBO} = \mathbb{E}_{p(n)}[\mathbb{E}_{q(z|n)}[\log(p(x|z)]] - \mathbb{E}_{p(n)}[KL[q(z|n)||p(z)]] \tag{2}$$

We define the loss over frames $\mathcal{L}_{frames}$ as the first of the two terms above. In the PS-VAE model, there are two inputs: frames $x$ and labels $y$. Therefore, in Equation (2), instead of writing the input likelihood as $p(x|z)$, we can now write it as $p(x, y|z)$. A simplifying assumption is made that $x$ and $y$ are conditionally independent given $z$, and thus we can directly write $\mathcal{L}_{frames+labels}$ as $\mathcal{L}_{frames} + \mathcal{L}_{labels}$, where $\mathcal{L}_{labels}$ is calculated by replacing $x$ with $y$ in $\mathcal{L}_{frames}$.

After assuming the prior $p(z)$ has a factorized form: $p(z) = \prod_i p(z_i)$, the KL term $\mathcal{L}_{KL}$ can be split as the addition of $\ell_{KL-s}$ and $\ell_{KL-u}$, i.e., the KL terms for the supervised and unsupervised latents, respectively. We decompose the KL term for the unsupervised latent as the following [2].

$$\begin{aligned}\mathcal{L}_{KL-u} &= \mathcal{L}_{ICMI} + \mathcal{L}_{TC} + \mathcal{L}_{DWKL} \\ &= KL[q(z_u, n)||q(z_u)p(n)] + KL[q(z_u)||\prod_j(z_{u,j})] + KL[q(z_{u,j})||\prod_j(z_{u,j})]\end{aligned} \tag{3}$$

where $j$ represents the latent dimension, $\mathcal{L}_{ICMI}$ is the index-code mutual information, which measures how well the latent encodes the corresponding input data. The term TC is short for total correlation, which measures the interdependency of each latent dimension. The third term, $\mathcal{L}_{DWKL}$ is the dimension-wise KL, which calculates the KL divergence for each dimension individually. Finally, the resulting subspace is forced to be orthogonal by applying orthogonal weights across all the different latents.

Table 1: Hyperparameter for different dataset

| Dataset | $\alpha$ | $\beta$ | $\sigma$ | $\gamma$ |
|---|---|---|---|---|
| Various contrast | 1000 | 5 | 5 | 500 |
| Multi-subject | 1000 | 5 | 15 | 500 |
| Social behavior | 1200 | $N/A$ | 20 | 200 |

Table 2: Latent dimensions and the prior distribution for different dataset

| Dataset | supervised | unsupervised | constrained | prior distribution |
|---|---|---|---|---|
| Various contrast | 5 | 2 | 3 | Swiss roll |
| Multi-subject | 5 | 2 | 2 | circle |
| Social behavior | 2 | 0 | 3 | hollow cylinder |

### D Model Architecture and Training

Our computational experiments were carried out using TensorFlow and Keras. The image decoder we use is symmetric to the encoder, with both of them containing 14 convolution layers. We applied the Adam optimizer with learning rate as $10^{-4}$. For the multi-subject dataset, we fixed our batch size to be 256 and trained for 50 epochs. For the freely-moving social behavior dataset, we trained for 500 epochs with batch size 128.

### E Choice of Hyperparameters

In the multi-subject dataset, four coefficients need to be decided for the objective function as indicated above: $\{\alpha, \beta, \sigma, \gamma\}$. There is a balance between the choice of $\beta$ and $\gamma$: properly choosing the values could separate the latent in the unsupervised space and the latents in both unsupervised and background space as well. A large separation of the background latent may potentially lead to unsatisfactory reconstruction results. The choice of kernel size $\sigma$ depends on the dataset, and should be larger than the number of distinct groups in our dataset; since in our current experiments, we have at most four groups, we set $\sigma = 15$. Moreover, we set $\alpha$ to 1000, $\beta$ to 5, $\gamma$ to 500. We set the dimensionality of the supervised latent space equal to the number of tracked video parts, which is 5 in our case. We set the dimensionality of the unsupervised latent space as 2, while that of the background latent space as 2.

In the social behavior task, we track the nose location as the supervised latent, since the other labels do not have a high variance (due to the alignment process). Additionally, we do not need any unsupervised latents to explain the individual's behavior. The CS latent in this setting has 3 dimensions. Here, $\alpha$ is 1200, $\gamma$ is 200, and the kernel size is 20.

The hyperparameters chosen for all three datasets are shown in Tables 1 and 2.

### F Motif Generation

A switching linear dynamical system (SLDS) consists of discrete latent state $z_t \in \{1, 2, ..K\}$, continuous latent state $x_t \in \mathbb{R}^M$, and the observation state $y_t \in \mathbb{R}^N$. Here, $t = 1, 2, 3, .., T$ is the time step, $T$ is the length of the input signal; $K$ is the number of discrete states; $M$ is the number of latent dimensions; $N$ is the observation dimensions. The discrete latent state $z_t$ follows the Markovian dynamics with the state transition matrix expressed as:

$$Q_{i,j} = P(z_t = j | z_{t-1} = i) \tag{5}$$

The continuous latent state $x_t$ has the following linear dynamical relations that determined by $z_t$.

$$x_{t+1} = A_{z_{t+1}} x_t + V_{z_{t+1}} u_t + b_{z_{t+1}} + w_t \tag{6}$$

Here, $A_{z_{t+1}}$ is the dynamic matrix at state $z_{t+1}$; $u_t$ is the input at time t, with $V_{z_{t+1}}$ being the control matrix; $b_{z_{t+1}}$ is the offset vector and $w_t$ being the noise which is generally the zero mean Gaussian. Here, our

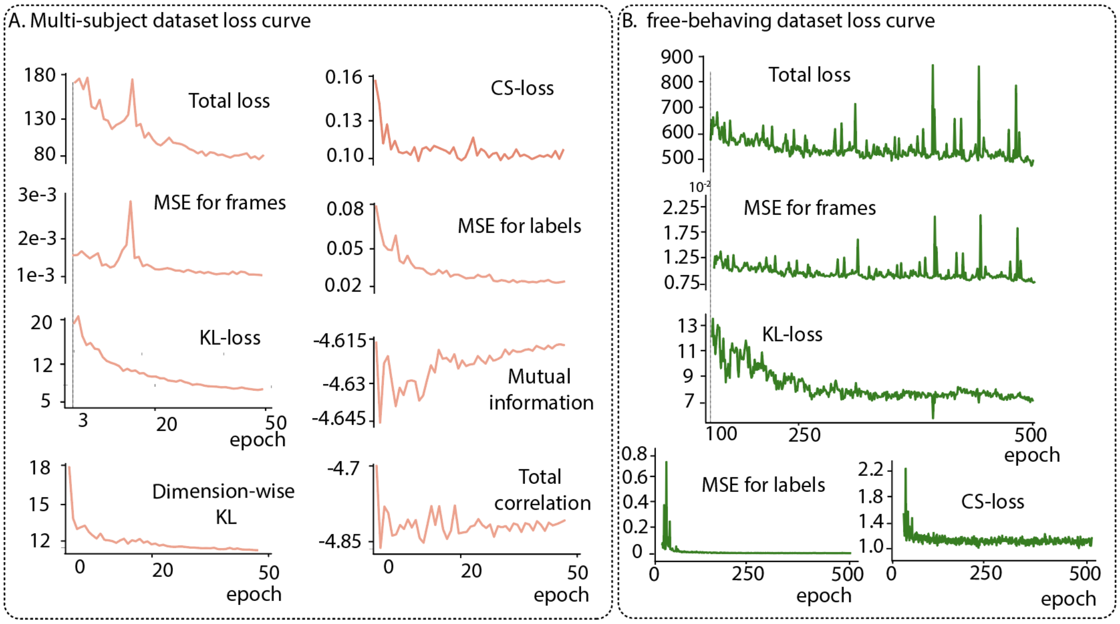


Figure 1: Loss curve for A. training the multi-subject dataset B. training the freely behaving dataset with the specified hyperparameters as in Tables 1 and 2.

observation model is in Gaussian case; therefore, the observation $y_t$ is expressed as:

$$y_t = C_{z_t} x_t + F_{z_t} u_t + d_{z_t} + v_t \tag{7}$$

Here, $C_{z_t}$ is the measurement matrix at state $z_t$; $F_{z_t}$ is the feedthrough matrix which directly feed the input into the observation; $d_{z_t}$ is the offset vector and $v_t$ is the noise. Here the update was accomplished by the Expectation-Maximization(EM) algorithm. In the E-step, the model updates the hyperparameters. In the M-step, the log-likelihood in Eq.7 is being maximized.

To implement the SLDS, we adopted the open source software from Linderman et al.[3]. We fit the SLDS using different latent dimensions, where the observation dimension was the order of latent dimension and the number of states was determined by visualizing the videos. We use SLDS's to model the motifs in the multi-subject dataset since the behaviors are well separated using their dynamics. We use K-means to model the motifs in the freely-moving social behavior dataset since the behaviors are well separated directly in state space. An autoregressive HMM (a simpler model than an SLDS) applied to the CS latents in the social behavior dataset leads to similar results as the K-means.

#### G Loss Curves

We show the learning curve for each loss term for both dataset to precisely quantify the model, in Fig. 1. For the multi-sujbect dataset (Fig. 1A), for the unsupervised latents, the final loss for dimension-wise KL, total correlation, and the mutual information are 11.7, $-4.8$, and $-4.6$, respectively. The final KL loss for the supervised latents is 5.06 and the final CSD loss for the CS latents is 0.1. For the free behaving dataset, the loss curves for each loss term are shown in Fig. 1B. By the end of the training process, the KL loss for the supervised latents is 7.01 and the CSD loss for the CS latents is 1.15.

#### H SVM

To further quantify the separation of the latents between different subjects, we applied a supervised classification method to decode the identity of the subject using each latent.

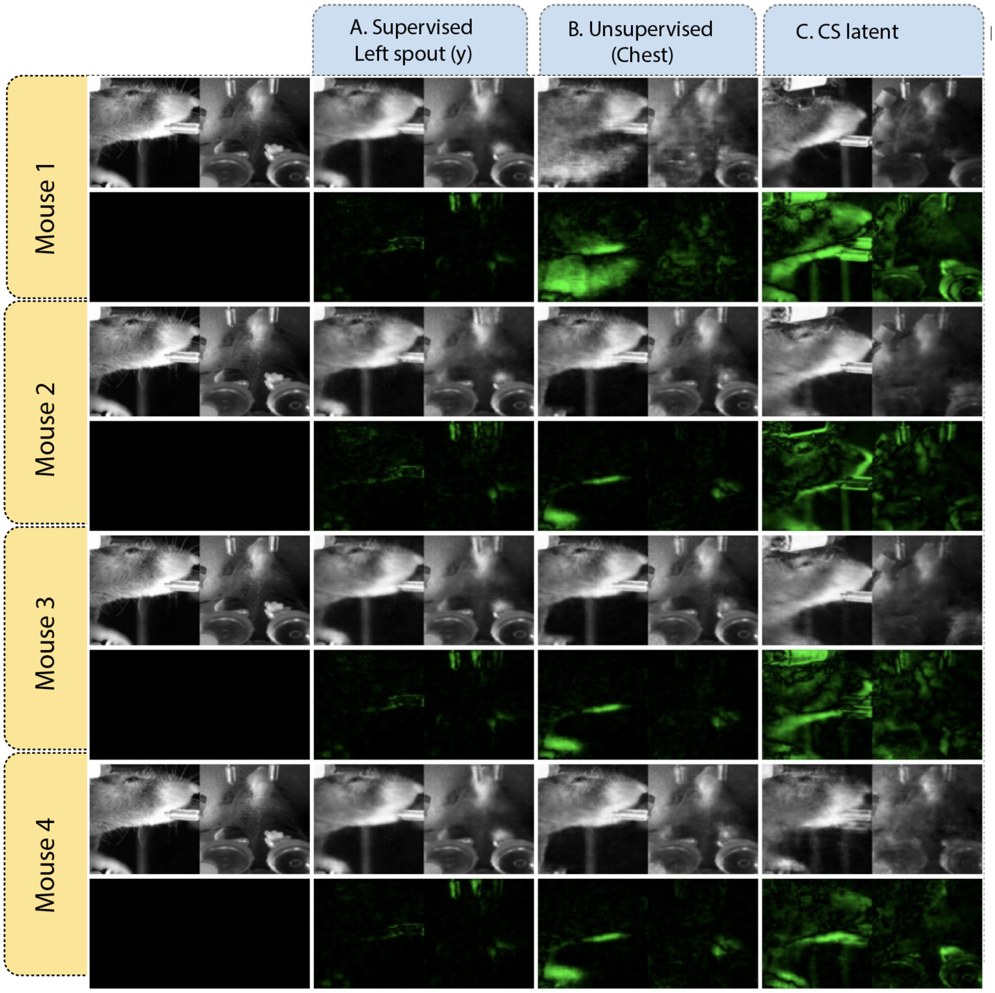


Figure 2: Latent traversals for the multi-subject dataset for the four mice with the same base image A. an example supervised latent, B. an example unsupervised latent, and C. an example CS latent. We see that the same base image (Mouse 3) is transformed into a different mouse each time when changing the CS latent.

After randomly shuffling all the latents, we split all the trials into training trials and test trials, with each mouse having 368 trials in the training set and 20 trials in the test set, and repeated this 5 times with different random seeds.

### I Latent traversal

For the multi-subject dataset, we tested the latent traversal with the same base image to validate the results, shown in Figure 2. Here, we randomly chose a frame from a mouse and changed each individual latent within different ranges as detailed in the Methods. For example, in Figure 2, the first row contains the output when the corresponding latent is changed to take on the maximum value from the range of Mouse 1. As in the figures in the main text, the upper images are the latent traversal images while the lower ones are the difference between the upper image and the original image. We see that the base image from Mouse 3 can be flexibly changed to produce a different mouse when changing the CS latent. Moreover, when changing the supervised and unsupervised latents for the different mice, Mouse 3 seems to be flexibly changing with these latents from different mice.

To better visualize the specialization of each latent, we generated the latent traversal videos for each latent with different base images. For different mouse, we first of all find the maximum and the minimum value for the specific latent. Then, change the latent within that range with 0.5 per step. Finally, concatenate all the latent traversal images into videos. The videos can be found in Supplementary Material 3.

We performed a similar visualization on the freely-moving social behavior dataset for the CS latents. The latent traversal videos can be found in Supplementary Material 4, and some clips from the videos are shown in Fig 3.

#### J Neural decoding models

The trials were first shuffled and then split into training and testing. Next, we employed the CS-VAE generated latent, and choose one example subject to decoded the behavior at time $t$ using the neural activity recorded between $t - 0.15$s and $t$. We applied four types of model to compare the performance. A linear model which directly map the neural activities into the behavior. A multilayer perceptron (MLP) with three dense layers to train the decoder. We used the Adam optimizer with learning rate decay from 0.1 with 0.3 decay rate for every 5 step. The batch size was fixed to be 150 and trained for 200 epochs. A LSTM model, which begin with a dense layer followed by a LSTM layer with drop out rate being 0.5 and another dense layer at the end. We applied the same training strategy as in MLP model.

We introduced a model based on transfer learning to perform the decoding test on the previously tested subject. The rest of the three mice were the input to the original training model. The procedures were similar as before, after the trials were being shuffled and split, we decoded the behavior directly with the raw neural activities with the time window being $0.15s$. After that, we implemented three perceptron layers for each of the three mice before the output of which went into a recurrent neural network (RNN). The RNN consisted of one long short-term memory (LSTM) layer with unit number of 64 and a drop out layer with rate being 0.5. We applied the Adam optimizer with learning rate decay from 0.1 with 0.3 decay rate for every 5 steps. The batch size was 150 and we trained for 200 epochs. After we finished training the original network, we transferred the RNN model to the new model which was applied to train for the fourth mouse alone. For the fourth mouse, the trials were split with different training and testing ratio. After applying the same steps to the data, the neural activities then went through a new perceptron layer before went through the pre-trained RNN model. We applied the Adam optimizer with the same learning rate decay procedures as well. We again, trained for 200 epochs with batch size being 128 this time. The trade-off between accuracy and time for different models can be found in Tables 3 and 4.

#### K Code

The code for training the CS-VAE can be found in Supplementary Material 7. The code can be executed by simply compiling the script 'train.py'.

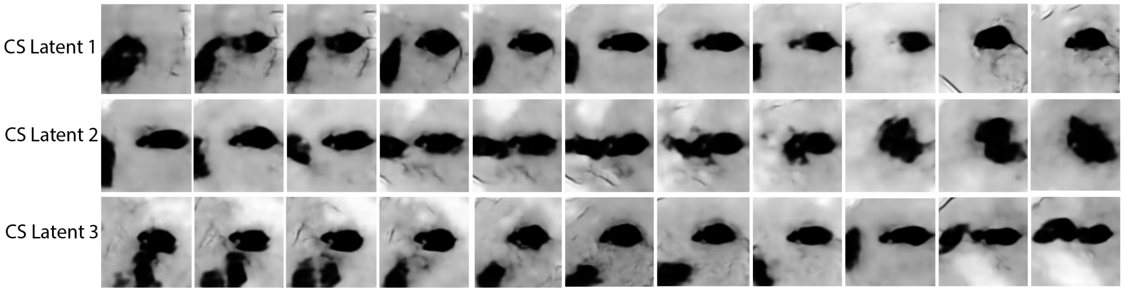


Figure 3: Latent traversals on the CS-latents for the freely-moving social behavior dataset. We see that the latents all encode for social interactions between the two mice.

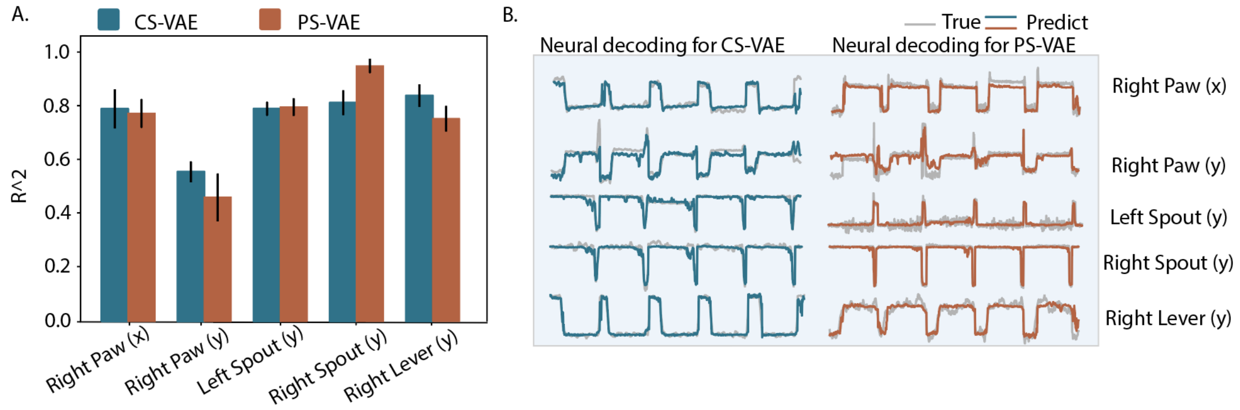


Figure 4: Neural decoding for CS-VAE vs. PS-VAE.

Table 3: Training size vs $R^2$ value for multi-subject dataset

| Training size | Linear model | Dense model | LSTM model | Transfer learning model |
|---|---|---|---|---|
| 67712 | $0.476 \pm 0.048$ | $0.580 \pm 0.058$ | $0.610 \pm 0.054$ | $0.590 \pm 0.050$ |
| 58512 | $0.478 \pm 0.014$ | $0.560 \pm 0.023$ | $0.595 \pm 0.022$ | $0.579 \pm 0.019$ |
| 49312 | $0.483 \pm 0.009$ | $0.556 \pm 0.014$ | $0.593 \pm 0.013$ | $0.576 \pm 0.011$ |
| 40112 | $0.476 \pm 0.013$ | $0.543 \pm 0.019$ | $0.576 \pm 0.019$ | $0.562 \pm 0.015$ |
| 30912 | $0.470 \pm 0.0.011$ | $0.529 \pm 0.015$ | $0.559 \pm 0.018$ | $0.552 \pm 0.013$ |
| 21712 | $0.458 \pm 0.010$ | $0.496 \pm 0.016$ | $0.524 \pm 0.017$ | $0.522 \pm 0.013$ |
| 12512 | $0.424 \pm 0.012$ | $0.461 \pm 0.0.019$ | $0.480 \pm 0.025$ | $0.485 \pm 0.018$ |
| 3312 | $0.269 \pm 0.030$ | $0.321 \pm 0.048$ | $0.325 \pm 0.057$ | $0.345 \pm 0.043$ |

[2] Whiteway, M. R. et al. Partitioning variability in animal behavioral videos using semi-supervised variational autoencoders. bioRxiv (2021).

[3] Linderman, S. et al. Bayesian Learning and Inference in Re- current Switching Linear Dynamical Systems. In Singh, A. Zhu, J. (eds.) Proceedings of the 20th International Conference on Artificial Intelligence and Statistics, vol. 54 of Proceedings of Machine Learning Research, 914–922 (PMLR, 2017).

Table 4: Training size vs time usage for multi-subject dataset

| Training size | Linear model | Dense model | LSTM model | Transfer learning model |
|---|---|---|---|---|
| 67712 | $1.442 \pm 0.282$ | $115.946 \pm 1.559$ | $169.801 \pm 5.961$ | $169.482 \pm 5.041$ |
| 58512 | $1.130 \pm 0.212$ | $80.428 \pm 1.586$ | $146.734 \pm 5.063$ | $151.771 \pm 4.162$ |
| 49312 | $0.937 \pm 0.194$ | $68.879 \pm 1.257$ | $122.500 \pm 2.283$ | $125.240 \pm 4.479$ |
| 40112 | $0.679 \pm 0.114$ | $56.449 \pm 1.119$ | $100.336 \pm 2.212$ | $102.923 \pm 3.391$ |
| 30912 | $0.484 \pm 0.079$ | $44.427 \pm 0.808$ | $78.850 \pm 1.771$ | $79.907 \pm 2.608$ |
| 21712 | $0.309 \pm 0.050$ | $31.968 \pm 0.574$ | $57.670 \pm 1.681$ | $56.032 \pm 1.549$ |
| 12512 | $0.162 \pm 0.008$ | $19.573 \pm 0.369$ | $35.557 \pm 1.050$ | $33.409 \pm 0.741$ |
| 3312 | $0.104 \pm 0.034$ | $7.292 \pm 0.092$ | $13.318 \pm 0.342$ | $11.365 \pm 0.336$ |
